# YBX3 overexpression in mesothelioma drives aberrant cell proliferation

**DOI:** 10.64898/2026.02.11.705262

**Authors:** Angela Rubio, Robert F Harvey, Andrew Craxton, Mark Southwood, Catherine Ficken, Catarina Franco, Lajos Kalmar, Goretti Muñoz, Ian Powley, Stephan Kamrad, Rui Guan, David Fernandez-Antoran, Kiran R Patil, John PC Le Quesne, Marion MacFarlane, Anne E Willis

## Abstract

Malignant pleural mesothelioma (MpM) is a lethal tumour closely linked to asbestos exposure and is a cancer of unmet clinical need with no known oncogenic drivers. Recent advancements in technologies to identify RNA-binding proteins (RBPs) has uncovered an emerging role for RBP-RNA interactions in cancer progression and we therefore assessed changes in the RBPome of patient-derived MpM cell lines. We identify over 350 RBPs showing altered RNA binding, with functions consistent with key cancer hallmarks, and discovered YBX3 as a potential oncoprotein driving cell proliferation in MpM. Mechanistically we show the impact of YBX3 on cell growth is achieved through its control of the expression of the amino acid transporter SLC7A5/LAT1, with increased amino acid uptake increasing protein synthesis rates. Notably, we show the inhibition of cell growth by YBX3 deletion is recapitulated by the clinically-relevant SLC7A5/LAT1 inhibitor JPH203. Finally, we demonstrate that JPH203 sensitizes MpM cells to radiotherapy, which could provide a promising therapeutic strategy for MpM.

**Teaser:** Higher levels of YBX3 expression in mesothelioma increases cell proliferation and protein synthesis rates by upregulation of amino acid uptake.

## INTRODUCTION

Malignant pleural mesothelioma (MpM), a lethal tumour originating from the mesothelial cell lining of the pleura, is closely linked to asbestos exposure (*1*). MpM is the most prevalent type of mesothelioma accounting for over 90,000 deaths per year globally, has a long latency period (often 40 years after the primary exposure), and the median overall survival following diagnosis is 12-18 months. Recent clinical trials have shown some improvements in survival, with immune checkpoint blockade using nivolumab plus ipilimumab outperforming chemotherapy with platinum-pemetrexed (*2*). However, despite these improvements, poor patient outcomes means that mesothelioma remains a cancer of unmet clinical need. Genomic analyses indicate that MpM is predominantly linked to the inactivation of tumour suppressor genes rather than mutations in targetable oncogenes (*3*, *4*). Notably, in 80–90% of MpM cases p16INK4a, p14ARF, NF2 and BAP1 are altered, and BAP1 and NF2 are mutated/deleted in ∼ 20% of cases (*3*, *5*). Dysregulated cytoplasmic control of gene expression also makes a major contribution to cancer development and progression (*6*), although this process has been little studied in mesothelioma. We have shown that in mesothelioma, enhanced signalling through mTORC1/2 results in a selective increase in the translation of mRNAs encoding proteins required for ribosome assembly. These alterations accelerate growth and drive disease progression. Importantly, we have shown that inhibition of mRNA translation, particularly through combined pharmacological targeting of mTORC1 and 2, reverses these changes and inhibits malignant cell growth *in vitro* and in *ex-vivo* tumour tissue from patients with end-stage disease (*7*).

Protein synthesis is controlled by RNA binding proteins (RBPs) that regulate the recruitment of RNAs to the ribosome, RNA localization and RNA turnover (*8*). Given these roles, it is therefore unsurprising that aberrant expression/activity of RBPs is associated with carcinogenesis (*9*), and poor patient outcome. The development of a range of technologies to decipher interactions between RBPs and their cognate RNAs, including RNA Interactome Capture (RIC) (*10*) and orthogonal organic phase separation (OOPS) (*11*), now allows for the interrogation of the cancer-associated RBPome. In this study, we used OOPS to characterize the RBPome changes in patient-derived primary MpM cell lines. We show that the RBPome is dysregulated in MpM and this promotes cell growth and disease progression. In particular, YBX3 was overexpressed in MpM and our data suggest that this protein acts as an oncogenic driver increasing the expression of the amino acid transporter SLC7A5/LAT1 (referred to as LAT1 from herein) which, in turn, increases amino acid uptake, to increase protein synthesis and promote proliferation. The impact of YBX3 depletion is recapitulated by a clinically-relevant LAT1 inhibitor, which sensitizes MpM derived cells to radiotherapy, highlighting a new more directed therapeutic option for this disease.

## RESULTS

### The RBPome is altered in MpM

Orthogonal organic phase separation (OOPS) uses acidic guanidinium thiocyanate-phenol-chloroform (AGPC) phase partition, whereby RNA migrates to the upper aqueous phase and proteins occupy the lower organic phase. UV cross-linking at 254 nm generates RNA–protein complexes that combine the physicochemical properties of the two molecules which then migrate to the aqueous–organic interface (Fig. 1A). OOPS enables the recovery of RBPs that bind all RNA species, including non-polyadenylated mRNAs, using less starting material than other methods (*11*). Control experiments showed that, as expected, the presence of proteins in the organic phase was dependent on UV crosslinking (RBPs, +/- CL) (Fig. S1A). OOPS was then applied to three MpM patient-derived cell lines (7T, 8T and 13T) (*12*) and healthy primary mesothelial control cells under standard growing conditions. The samples were then TMT labelled and analysed by mass spectrometry. After computational analysis, 1661 potential RBPs were identified (Fig. 1B), of which 1246 (75%) were either identified by OOPS in other cell lines (*11*) or annotated with the GO term RBP (Fig. 1C). PCA plots show that healthy mesothelial cells and MpM-derived cells group along the first component for both sample types, total and potential RBPs (Fig. S1B).

**Fig. 1.**
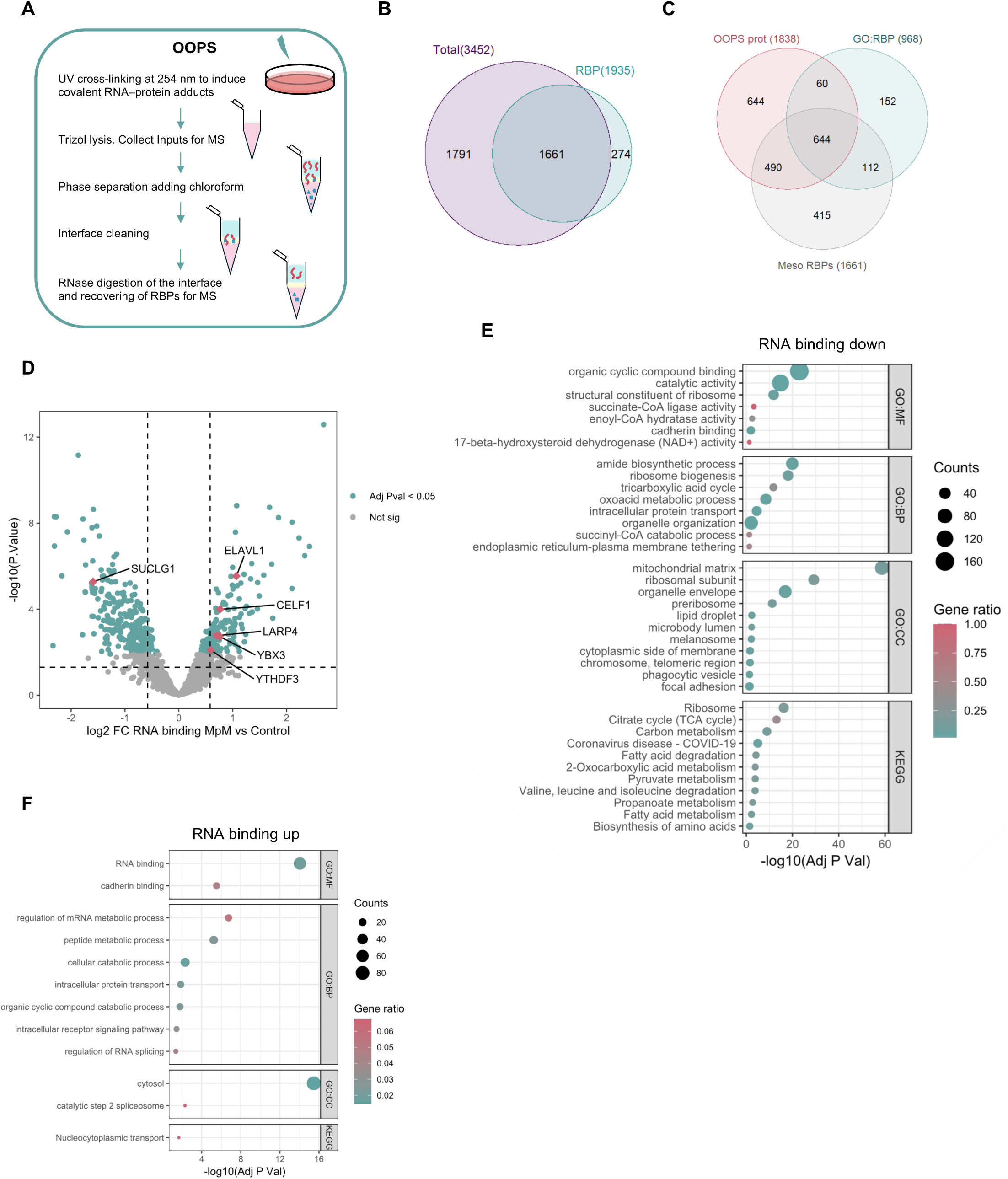
The RBPome is dysregulated in MpM. (**A**). Schematic representation of the OOPS protocol. Cells are cross-linked to induce RNA-protein adducts, which migrate to interface in a Trizol/chloroform extraction. RNase digestion and a further AGPC separation release RBPs in the organic phase. (**B**) Venn diagram comparing the number of proteins identified in the input samples (Total) and interface samples (RBPs). (**C**) Venn diagram comparing the number of intersecting MpM RBPs from (B) compared with the number of potential RBPs detected by OOPS in other cell lines (*11*) and the number of proteins annotated in the GO term RBP. (**D**) Volcano plot comparing RBPs from OOPS mass spectrometry data. Adjusted P value < 5*10^−2^ and FC > 1.5 was considered significant in RBP versus Total samples. RBPs mentioned in the text are highlighted in pink. The vertical dotted lines correspond to 1.5-fold differences and the horizontal dotted lines to a P. Value of 0.05. OOPS data are provided in Supplementary Table 1. (**E** and **F**) Dot plots depicting GO and KEGG functional enrichment analysis of RBPs showing (E) a decrease or (F) an increase in their RNA-binding.

Over 350 RBPs displayed significant changes in RNA association, i.e. those below a threshold of 5x10^−2^ for the adjusted P value and a cut-off of 1.5-fold minimal change in RBP versus total samples (Table 1, Table 2 and Supplementary Table 1), and changes in RNA binding also correlated with total expression (Fig. S1C, D). Many of the RBPs which showed a significant increase in MpM cell lines including ELAVL1, CELF1, YBX3 or LARP4 (Fig.1D, S1C group n=47), have a role in cancer development and metastasis. For example, CELF1 promotes alternative splicing of several pre-mRNAs involved in tumorigenesis and is overexpressed in both oral and colon cancers (*13–15*). Similarly, an increase of the cytoplasmic abundance of ELAVL1 (HuR) is commonly associated with poor clinical outcomes and altered expression in multiple tumour types, including pancreatic, colon, breast, prostate, ovarian, and MpM (*16–22*). Aberrant expression of YBX3, a DNA/RNA binding protein involved in gene transcription, translation and mRNA stability (*23*, *24*), has been associated with tumour growth and invasion in colorectal cancer (*25*), gastric cancer cells (*26*) and nasopharyngeal carcinoma (*26*). In addition, we observed a change in the RNA-binding of LARP4, which has been implicated in the inhibition of migration and invasion of prostate and breast cancer cells through regulation of mRNA translation by interaction with poly(A)-binding protein (PABP) (*27*). We also identified significant changes in the RNA-binding of the m6A reader YTHDF3 (Fig. S1C, D group n=58), and alterations in the expression of this protein and its co-regulators have also been associated with tumorigenesis (*28–30*). Functional enrichment analysis of RBPs that showed decreased RNA binding (Fig. 1E) identified enzymes of the tricarboxylic acid cycle (TCA), RBPs localised in the mitochondrial matrix and, overall, associated with metabolism. Conversely, RBPs with increased RNA binding showed enrichment of GO terms related to RNA binding, RNA metabolic processes and splicing, consistent with the metabolic reprogramming and dysregulated RNA splicing as hallmarks of cancer (Fig. 1F).

**Table 1.** Proteins from MpM cell lines versus control cells significantly up or downregulated (threshold of adjusted P value < 5*10^−2^ and FC > 1.5 were chosen), or without significant changes from OOPS experiment. RBP comes from the interface, Total from the inputs and Diff integrates both interface and inputs changes.

|  | Mesothelioma vs<br>Control RBP | Mesothelioma vs<br>Control Total | Diff Mesothelioma |
| --- | --- | --- | --- |
| Down | 345 | 275 | 226 |
| NotSig | 997 | 1093 | 1300 |
| Up | 319 | 293 | 135 |

**Table 2.** Proteins from each MpM cell line versus control significantly up or downregulated, or without significant changes from OOPS experiment. RBP comes from the interface, Total from the inputs and Diff integrates both interface and inputs changes.

|  | M7T vs<br>Control<br>RBP | M8T vs<br>Control<br>RBP | M13T vs<br>Control<br>RBP | M7T vs<br>Control<br>Total | M8T vs<br>Control<br>Total | M13T vs<br>Control<br>Total | Diff M7T | Diff M8T | Diff<br>M13T |
| --- | --- | --- | --- | --- | --- | --- | --- | --- | --- |
| Down | 330 | 348 | 394 | 244 | 315 | 314 | 173 | 140 | 274 |
| NotSig | 1026 | 1008 | 898 | 1169 | 994 | 1032 | 1367 | 1423 | 1210 |
| Up | 305 | 305 | 369 | 248 | 352 | 315 | 121 | 98 | 177 |

The total proteome of the three mesothelioma cell lines was also compared to the control primary mesothelial cells and 1292 proteins were either up or downregulated (Table 3 and Supplementary Table 2). The upregulated proteins group was enriched for proteins involved in DNA metabolic processes, such as replication or repair, RNA metabolism and localization, and mitotic cell cycle (e.g. CDK1, CDK2, MCM3, MCM6, PCNA, ENSA) (Fig S1E). The downregulated protein group has GO terms enriched for metabolic process, including oxidoreductase activity, lipid and amino acid metabolism. In addition, many proteins related to apoptosis were downregulated (e.g. CDK5, SOD2, PDCD4) (Fig S1F).

**Table 3.** Proteins from MpM cell lines versus control cells and each MpM cell line versus control cells significantly up or downregulated from the inputs.

|  | Mesothelioma vs<br>Control | M7T vs Control | M8T vs Control | M13T vs Control |
| --- | --- | --- | --- | --- |
| Down | 659 | 601 | 748 | 698 |
| NotSig | 2160 | 2323 | 1957 | 2055 |
| Up | 633 | 528 | 747 | 699 |

Overall, the upregulation of cell cycle and downregulation of cell death-related proteins correlates with a proteomic profile of malignant proliferating tumour cells, and is consistent with the hypothesis that changes in the MpM RBPome plays a central role in the tumour proliferation phenotype.

### YBX3 is overexpressed in MpM

Given that RBPs regulate all aspects of mRNA translation, and we have previously demonstrated reprogramming of the translatome in MpM (*7*), it was important to determine the RBPs that might regulate these processes. A crucial step for sustaining the rates of mRNA translation in tumour cells is the import of amino acids, which is mediated by transporters such as LAT1 and SLC3A2. Moreover, the RBP YBX3 has previously been shown to regulate the expression of LAT1 and SLC3A2 by binding to their mRNAs (*31*), and metabolic reprogramming in MpM has been described, suggesting a potential role for YBX3 in the pathogenesis of mesothelioma. Therefore, to investigate the role of YBX3 in MpM we first validated the mass spectrometry data for YBX3 RNA binding (Fig. 2A) using both OOPS and RNA Interactome Capture (RIC) by western blotting. Both techniques showed that YBX3 exhibits a greater interaction with RNA in MpM cells compared to control mesothelial cells and relative to DDX3, which did not exhibit significant changes in our OOPS dataset (Fig. 2B). Furthermore, total levels of YBX3 were analysed from three MpM-derived cell lines (7T, 8T, and 13T) using western blotting. These data demonstrate the upregulation of YBX3 in MpM compared to control mesothelial cells (Fig. 2C).

**Fig. 2.**
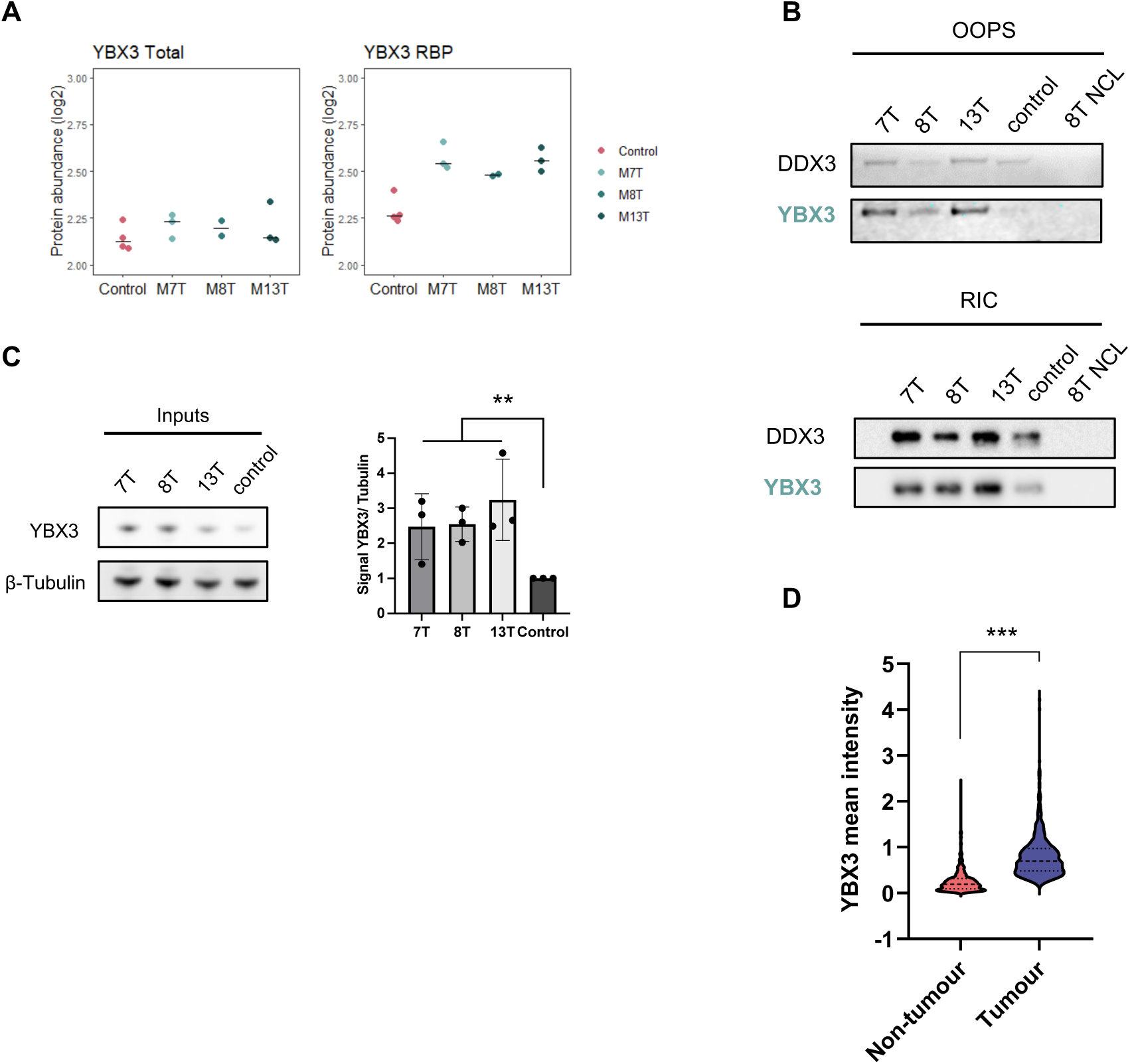
YBX3 is overexpressed in MpM. (**A**) Comparison of protein abundance from mass spectrometry data of the RBP YBX3 between control and MpM cell lines in Total and RBP samples. (**B**) Western blots comparing YBX3 levels in RBP samples from OOPS (above) and RIC (below) methods between MpM cell lines and control cells. 8T NCL is a non-crosslinked sample used as negative control for RNA binding. DDX3 was used as an invariable control RBP. (**C**) Representative western blot and quantification comparing YBX3 levels in inputs samples between MpM cell lines and control mesothelial cells. Error bars represent means ± SD (n = 3 independent experiments). Significance assed by unpaired Student’s t test (** = p<0.01). Tubulin was employed as a loading control. (**D**) Violin plot comparing YBX3 mean intensity from TMA immunohistochemical staining in tumour and non-tumour areas. Significance assed by unpaired Student’s t test (*** = p<0.001).

To ascertain whether YBX3 overexpression was prevalent during the development of MpM, we expanded our assessment of YBX3 expression to patient material present on a mesothelioma tissue microarray (TMA). This TMA comprises formalin-fixed surgical tumour samples from 512 individuals with mesothelioma. The TMAs were probed with antibodies against YBX3 and MNF116 (pan-cytokeratin marker) to assess the tumour versus non-tumour areas. Quantitative protein expression data were obtained from images using Visiopharm® software, and YBX3 mean intensity from TMA immunohistochemical staining was significantly increased in tumour areas (Fig. 2D and Fig. S2). Therefore, we conclude that YBX3 is overexpressed in MpM (tissue and cell lines) and the increase in the RNA-binding of YBX3 observed by OOPS is likely caused by increased protein expression in MpM and increased interactions with target mRNA. We next wanted to determine if the overexpression of YBX3 was an early event during MpM development. To answer this question we utilised mouse tissue from a previous longitudinal study of fibre-induced MpM development (*32*) and carried out immunohistochemical analysis of YBX3 expression. Whereas YBX3 expression was minimal in the vehicle control, YBX3 was upregulated in mice that developed pleural tumours that contained abundant mesothelin-positive cells, correlating with our human TMA data (Fig. S3A - C). Interestingly, enhanced YBX3 expression was also observed in mice that developed late-stage lesions (Fig. S3D).

Taken together, these data demonstrate that the overexpression of YBX3 in MpM and this results in the increase of YBX3 binding to its target RNAs.

### YBX3 depletion reduces MpM proliferation and protein synthesis rates

Since YBX3 has previously been shown to regulate cell proliferation and cancer progression in some tumour types (*26*, *33–36*), we addressed whether elevated YBX3 expression enhanced MpM proliferation. Small interfering RNA (siRNA) specific to YBX3 (siYBX3) were used to knockdown YBX3 in 8T and 13T MpM cell lines and cell growth was monitored using the IncuCyte Live-Cell system. These data show that YBX3 depletion significantly reduced proliferation of both 8T and 13T MpM cell lines over the course of 72 hours (Fig. 3A, 3B). To determine if reduced proliferation upon YBX3 depletion was accompanied by cell cycle arrest or cell death, cell cycle distribution was assessed using propidium iodide (PI) staining and cell death using annexin V and DRAQ7 staining (Fig. S4A and S4B). Importantly, there were no changes in cell cycle distribution or cell death upon YBX3 knockdown, suggesting that all stages of the cell cycle may be slowed in response to YBX3 depletion.

**Fig. 3.**
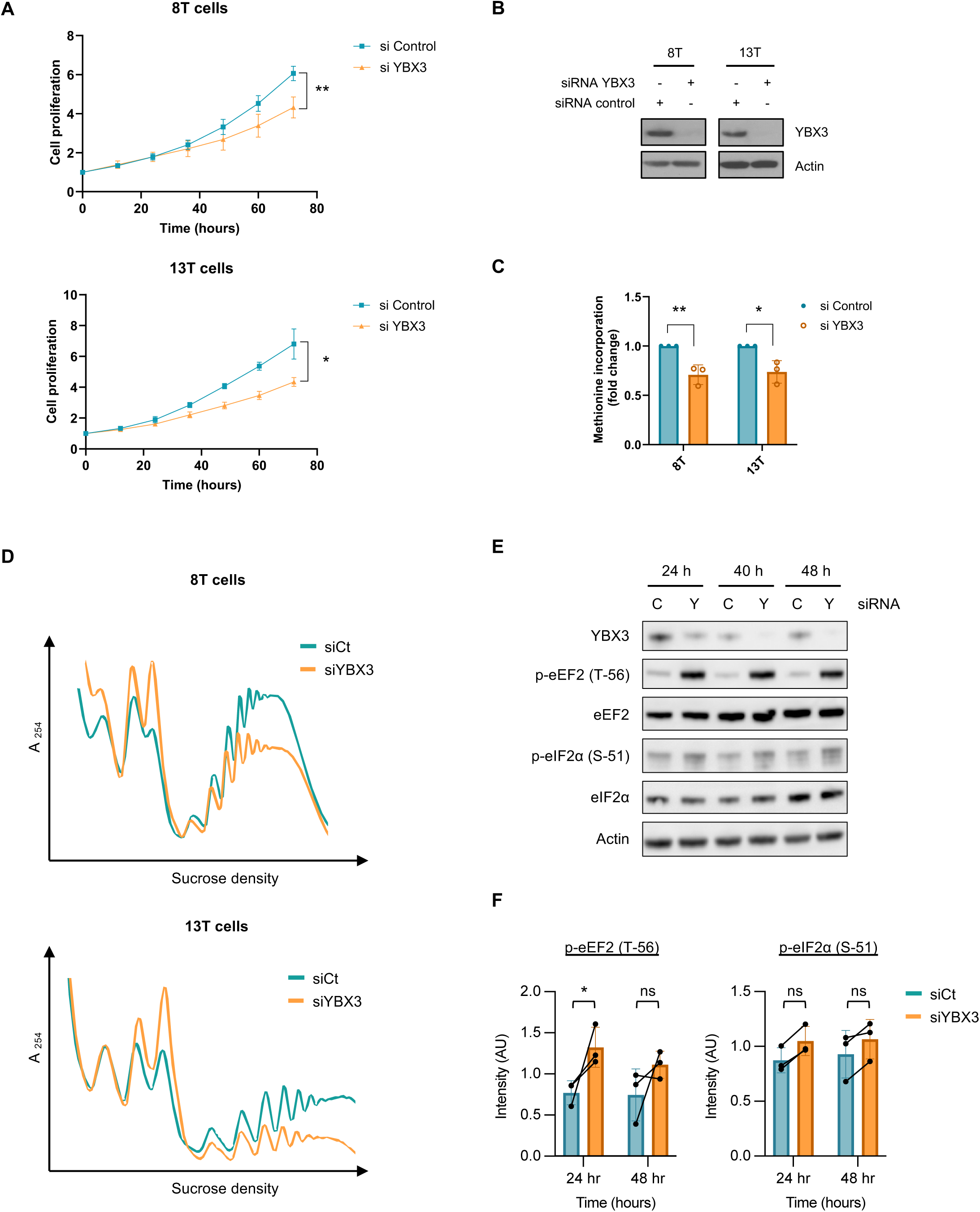
YBX3 promotes proliferation and protein synthesis in MpM. **(A)** Cell proliferation profiles from the IncuCyte Live-Cell instrument. MpM cells were treated for 24 hours with siRNA specific for YBX3 or non-targeting control before monitoring growth during the following 72 hours. Error bars represent means ± SD (n = 3 independent experiments). Significance assed by unpaired Student’s t test (* = p<0.05 and ** = p<0.01). (**B**) In parallel with (A), representative western blot of YBX3 protein expression levels confirming the knockdown efficiency at 72 hours. (**C**) MpM cell lines 8T and 13T cells transfected with siRNA specific for YBX3 or non-targeting control were pulse-labelled with [35S]-methionine for 30 min. Total counts per minute were normalized to total protein, and values are shown as a fold change for each cell line. Error bars represent means ± SD (n = 3 independent experiments). Significance assed by unpaired Student’s t test (* = p<0.05 and ** = p<0.01). (**D**) Comparison of polysome profiles from MpM cell lines transfected with siRNA specific for YBX3 or non-targeting control for 72 hours. Cytoplasmic lysates were centrifuged at 38,000 rpm through 10 to 50% sucrose gradients at 4°C for 2 hours, and absorbance was measured at 254 nm using a flow rate of 1 ml/min. Profiles shown were representative of two independent experiments. (**E**) Representative western blots from 8T MpM cells treated with siRNA specific for YBX3 or non-targeting control for the indicated times. Cells were lysed and analysed by immunoblotting with the indicated antibodies. (**F**) Quantification of eEF2 (T-56) and eIF2α (S-51) phosphorylation from (F). Error bars represent means ± SD (n = 3 independent experiments). Significance assed by unpaired Student’s t test (* = p<0.05).

As YBX3 regulates intracellular amino acid levels through the stabilization of mRNAs encoding amino acid transporters (*31*), and that protein synthesis rates and cell proliferation are intricately linked (*37*), we next assessed the levels of protein synthesis in 8T and 13T MpM cell lines following YBX3 depletion. We first utilised a radiolabelled [^35^S]-methionine incorporation assay which showed a ∼30% reduction in translation initiation synthesis rates following depletion of YBX3 for 72 hours (Fig. 3C). Moreover, polysome profiling showed a decrease in polysomes (actively translating ribosomes) and an increase in the sub polysomes in 8T and 13T MpM cell lines (Fig. 3D), consistent with an inhibition of mRNA translation. Finally, to further understand the impact on translation we assessed the status of elongation and initiation by monitoring the phosphorylation of eEF2 and eIF2⍺ at earlier timepoints of YBX3 depletion. Interestingly, eEF2 phosphorylation was elevated upon YBX3 depletion (Fig 3E and 3F), and although only significant at 24 hours, the trend of enhanced eEF2 phosphorylation was also observed at 48 hours. In addition, we observed a small increase in eIF2⍺ phosphorylation (Fig 3E and 3F), which although not significant, correlates with the decreases in [^35^S]-methionine incorporation (Fig 3C).

Taken together, these data suggest that over-expression of YBX3 in MpM may promote mRNA translation to enhance cell proliferation.

### YBX3 binds LAT1 mRNA and enhances its expression

YBX3 stabilises the mRNAs encoding the amino acid transporters LAT1 and SLC3A2 by binding to their 3’ UTRs, establishing a direct molecular function of YBX3 in amino acid transport (*31*) and control of protein synthesis rates. To investigate whether YBX3 was also upstream of LAT1 and SLC3A2 in MpM, we monitored expression of these proteins following siRNA-mediated YBX3 depletion. These data suggest that LAT1 expression decreased when YBX3 was reduced, particularly in 8T MpM cells, whereas the effect on SLC3A2 was minimal (Fig. 4A, S5A). To assess the binding of YBX3 to LAT1 mRNA in MpM, ribonucleoprotein (RNP) complex immunoprecipitation and qPCR (RIP-qPCR) was carried out, and we observed an enrichment of YBX3 binding to LAT1 mRNA compared to the non-YBX3 bound mRNA SLC25A6 (Fig. 4B, S5B, S5C). Additionally, siRNA depletion of LAT1 resulted in a similar decrease of cell proliferation as we observed in response to YBX3 depletion (Fig. 4C, S5D), suggesting the upregulation of YBX3 may increase LAT1 expression in MpM by binding to LAT1 mRNA. Consistent with this hypothesis, LAT1 overexpression in mice that developed tumours or lesions correlated with YBX3 levels (Fig. S3), and LAT1 was significantly overexpressed in tumour areas from our human TMAs (Fig. 4D), with parallel staining for YBX3 showing a strong correlation between both proteins in all tumour subtypes (Fig. 4E, F, G, Table 4, Supplementary Table 4). Importantly, these data showed that patients with high protein expression of both YBX3 and LAT1 have a significantly worse overall survival rate (Fig. 4G, Supplementary Table 4), strongly suggesting that these proteins play a key role in MpM development and progression.

**Fig. 4.**
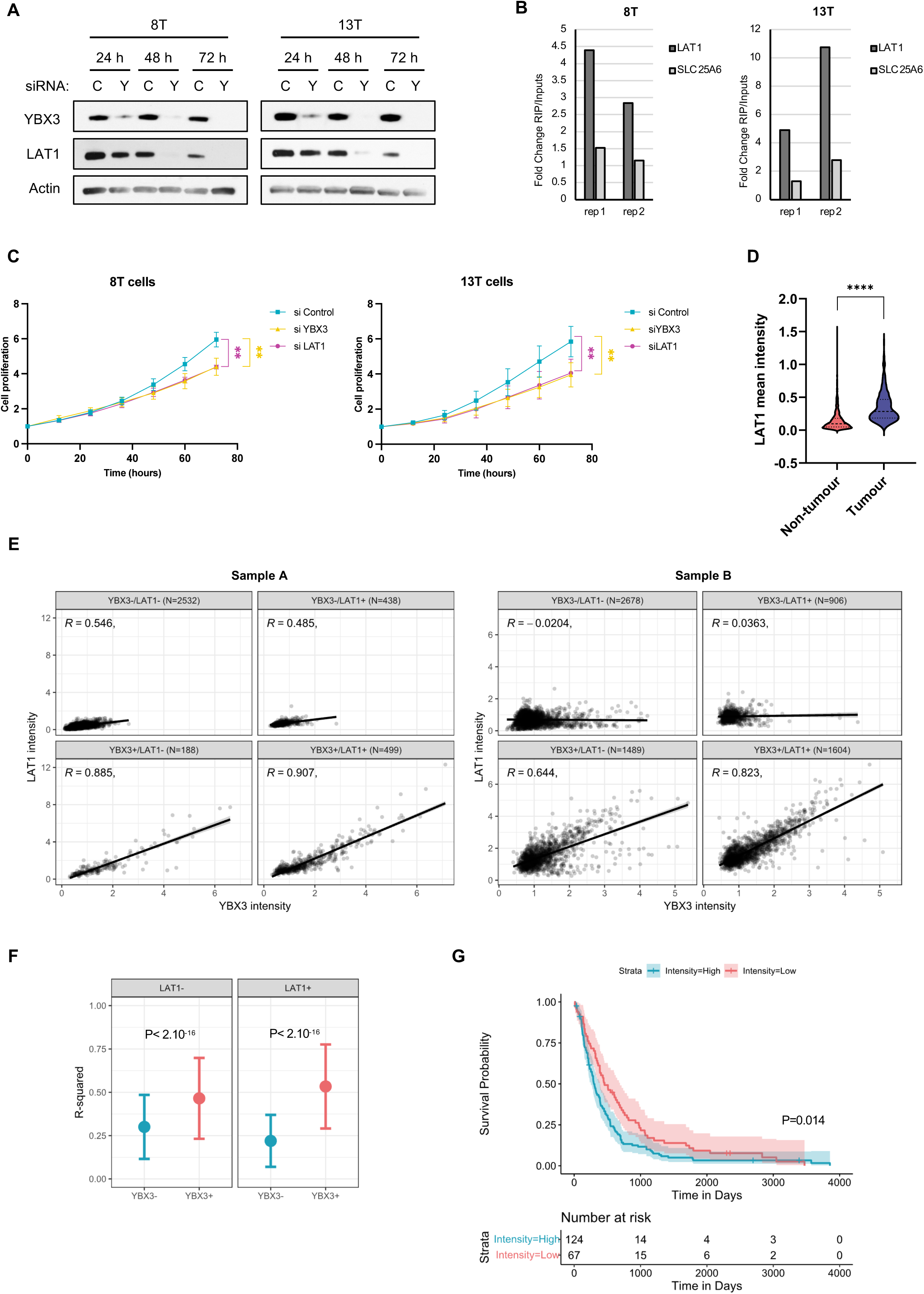
The effects of YBX3 on cell proliferation are mediated through LAT1. (**A**) MpM cells were treated with siRNA specific for YBX3 (Y) or non-targeting control (C) for the indicated times. Cells were lysed and analysed by immunoblotting with the indicated antibodies to show LAT1 expression decreases after YBX3 depletion. Representative blots of 3 independent experiments are shown. (**B**) YBX3 ribonucleoprotein (RNP) complex immunoprecipitation followed by LAT1 qPCR (RIP-qPCR) shows an enrichment of LAT1 mRNA binding compared to a non-YBX3 target control mRNA (SLC25A6). (**C**) Cell proliferation profiles from the IncuCyte Live-Cell instrument. MpM cells were treated for 24 hours with siRNA specific for YBX3, LAT1 or non-targeting control before monitoring growth during the following 72 hours. Error bars represent means ± SD (n = 3 independent experiments). Statistical analysis was carried out using one-way ANOVA with Tukey’s multiple comparisons test (** = p<0.01) (**D**) Violin plot comparing LAT1 mean intensity from TMA immunohistochemical staining in tumour and non-tumour areas. Significance assed by unpaired Student’s t test (**** = p<0.0001). (**E**) Scatter plots of YBX3 and LAT1 mean intensities at cell level from two TMA samples (sample A and sample B) showing correlations between expression. Pearson R and cell number by subset is shown. (**F**) Plot of R-squared means (dots) and SD (error bars) of YBX3 and LAT1 intensities at cell level by cell subsets. P value by unpaired Student’s t test is shown. (**G**) Kaplan–Meier survival curves with log-rank analysis of patients with high or low YBX3 and LAT1 mean intensities on the TMA.

**Table 4.** R-squared means of YBX3 and LAT1 intensities by mesothelioma subtype.

| Cell subset | Epithelioid | Sarcomatoid | Biphasic |
| --- | --- | --- | --- |
| SLC7A5- / YBX3- | 0.298 ± 0.179 | 0.321 ± 0.225 | 0.307 ± 0.202 |
| SLC7A5- / YBX3+ | 0.465 ± 0.236 | 0.465 ± 0.216 | 0.466 ± 0.224 |
| SLC7A5+ / YBX3- | 0.218 ± 0.149 | 0.217 ± 0.169 | 0.232 ± 0.151 |
| SLC7A5+ / YBX3+ | 0.53 ± 0.246 | 0.54 ± 0.244 | 0.55 ± 0.226 |

### Inhibition of LAT1-mediated amino acid transport reduces protein synthesis and MpM cell proliferation

As LAT1 imports large neutral amino acids (such as leucine, isoleucine, valine, phenylalanine, tyrosine, tryptophan, methionine, and histidine) across the plasma membrane (*38–40*), it was important to identify the amino acids transported by LAT1 in MpM. To address this question, the abundance of amino acids was measured in MpM 8T and 13T cell lines by targeted LC/MS after 3, 6 or 16 hours of exposure to the specific LAT1 inhibitor JPH203 (*41*). This analysis revealed that the LAT1 amino acid substrates phenylalanine, tyrosine or tryptophan were downregulated in both MpM cell lines and there was compensation of some non-essential amino acids and accumulation of glutamine (Fig. 5A). Methionine, which is also a LAT1 substrate, showed significantly increased intracellular levels and these data are consistent with previous reports on amino acid changes in LAT1 knockout in small intestinal tissues of mice (*42*). In addition, leucine levels were significantly decreased in 13T (Supplementary Table 3), correlating with previous investigations which showed that LAT1 is the primary transporter of leucine in some cancer cell lines (*43*, *44*). By contrast, arginine, a non LAT1 substrate amino acid relevant in cancer proliferation, showed significantly lower levels after JPH203 treatment in 8T cells (Supplementary Table 3), which together with differences in leucine changes suggests there are some variations between cell lines. While some subtle differences between our cell lines may be explained by the fact that they are patient-derived, these data reveal that JPH203 disrupts LAT1 substrate intake with accumulation of intracellular glutamine in MpM.

**Fig. 5.**
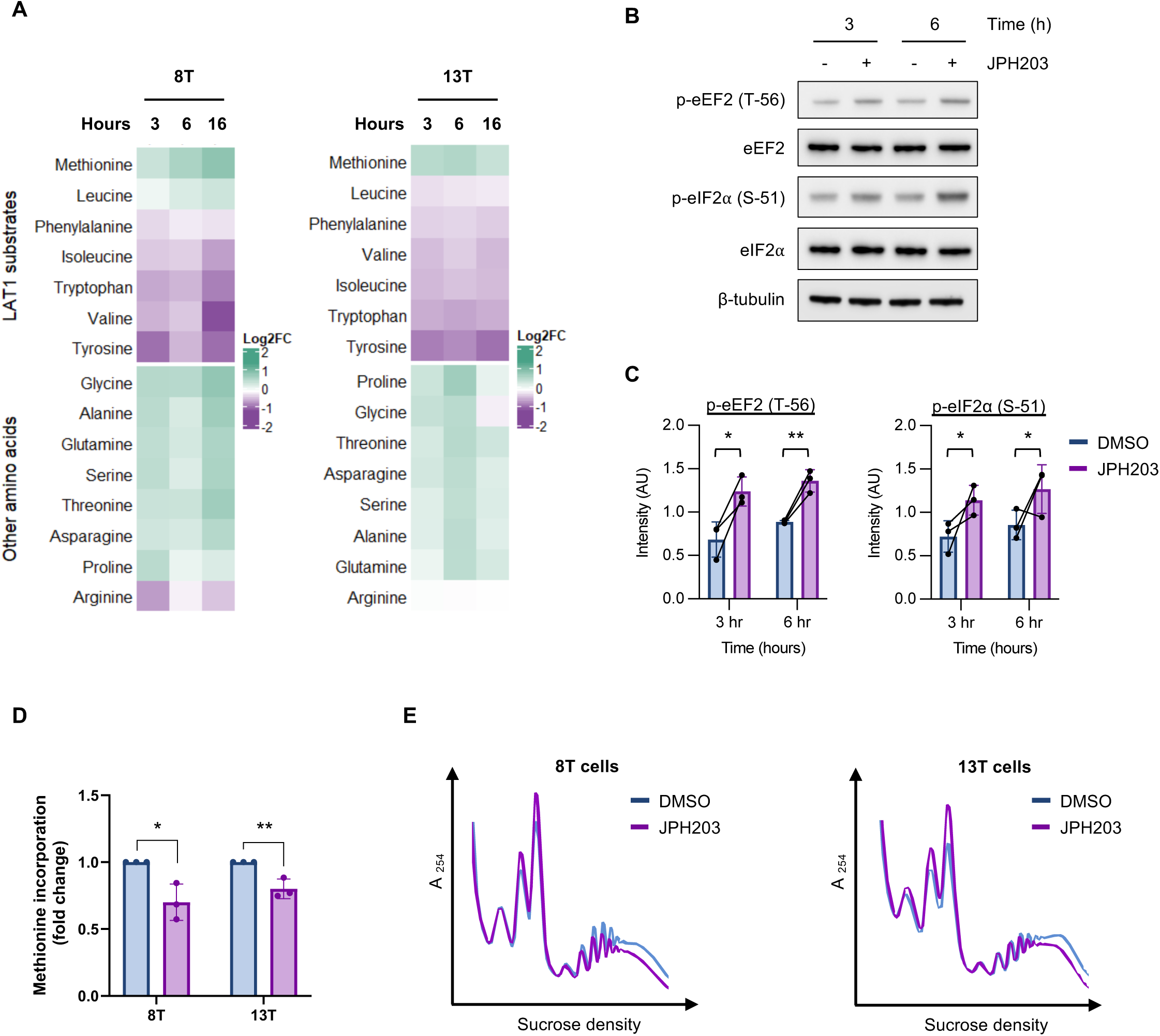
LAT1 inhibition affects amino acid transport, protein synthesis and proliferation in MpM. **(A)** Abundance of amino acids in 8T and 13T MpM cell lines treated with JPH203 (10 µM) for the indicated times using LC-MS. Heatmap shows mean relative abundance (n = 6 independent biological replicates per group). **(B)** Representative western blots from 8T cells were treated with JPH203 (10 µM) or DMSO for the indicated times. **(C)** Quantification of eEF2 (T-56) and eIF2α (S-51) phosphorylation from (B). Error bars represent means ± SD (n = 3 independent experiments). Significance assed by unpaired Student’s t test (* = p<0.05 and ** = p<0.01). **(D)** MpM cell lines 8T and 13T cells treated with JPH203 (10 µM) or DMSO for 6 hours were pulse-labelled with [35S]-methionine for 30 min. Total counts per minute were normalized to total protein and values are shown as a fold change for each cell line. Error bars represent means ± SD (n = 3 independent experiments) and significance assed by unpaired Student’s t test (* = p<0.05 and ** = p<0.01). **(E)** Comparison of polysome profiles from MpM 8T and 13T cell lines treated with JPH203 (10 µM) or DMSO for 6 hours. Cytoplasmic lysates were centrifuged at 38,000 rpm through 10 to 50% sucrose gradients at 4°C for 2 hours, and absorbance was measured at 254 nm using a flow rate of 1 ml/min. Profiles shown were representative of two independent experiments.

To determine the impact of amino acid imbalance induced by LAT1 inhibition on mRNA translation, we assessed the phosphorylation status of canonical initiation and elongation factors in MpM 8T cells by western blotting. Importantly, JPH203 treatment significantly increased the phosphorylation of eEF2 and eIF2α, indicators of elongation and initiation inhibition, respectively, within 3 hours (Fig. 5B and 5C). Moreover, treatment with JPH203 for 6 hours decreased global protein synthesis rates and reduced polysomes with a concurrent increase of sub polysomes (Fig. 5D and E) in both 8T and 13T MpM cells, consistent with previous observations following YBX3 depletion (Fig. 3C, 3D). The availability of amino acids is also sensed by rag GTPases which result in mTORC1 activation at the lysosome (*45*). Given that mTORC1/2 signalling has previously been shown to be upregulated in MpM (*7*), we wanted to determine if the amino acid imbalance induced by acute LAT1 inhibition inhibited mTORC1 signalling. Although we observed a subtle decrease in the phosphorylation of mTORC1 substrates 4EBP1 and p70 S6K following JPH203 treatment for 6 hours, neither were significant (Fig. S6). These data suggest that the acute response to LAT1 inhibition may be inhibiting mRNA translation by limiting amino acid availability to initiating and elongating ribosomes rather than mTORC1 activity.

Taken together, these data suggest that LAT1 inhibition by JPH203 decreases protein synthesis rates in MpM cell lines.

### JPH203 sensitise MpM cells to radiation by increasing senescence

At present, the standard of care for MpM is chemotherapy based on a combination of pemetrexed and cisplatin, although several immunotherapies are in clinical trials showing promising results (*46*). Conversely, hemithoracic radiation is only currently used in combination with surgery and or chemotherapy (*47*). As protein synthesis was downregulated upon treatment with JPH203 (Fig. 5), we next asked whether JPH203 could provide an alternative treatment option for MpM by determining the anti-proliferative effect in MpM cell lines. Importantly, treatment with JPH203 significantly reduced proliferation rates of both 8T and 13T MpM cell lines (Fig. 6A), recapitulating our previous observations following depletion of LAT1 (Fig. 4C).

**Fig. 6.**
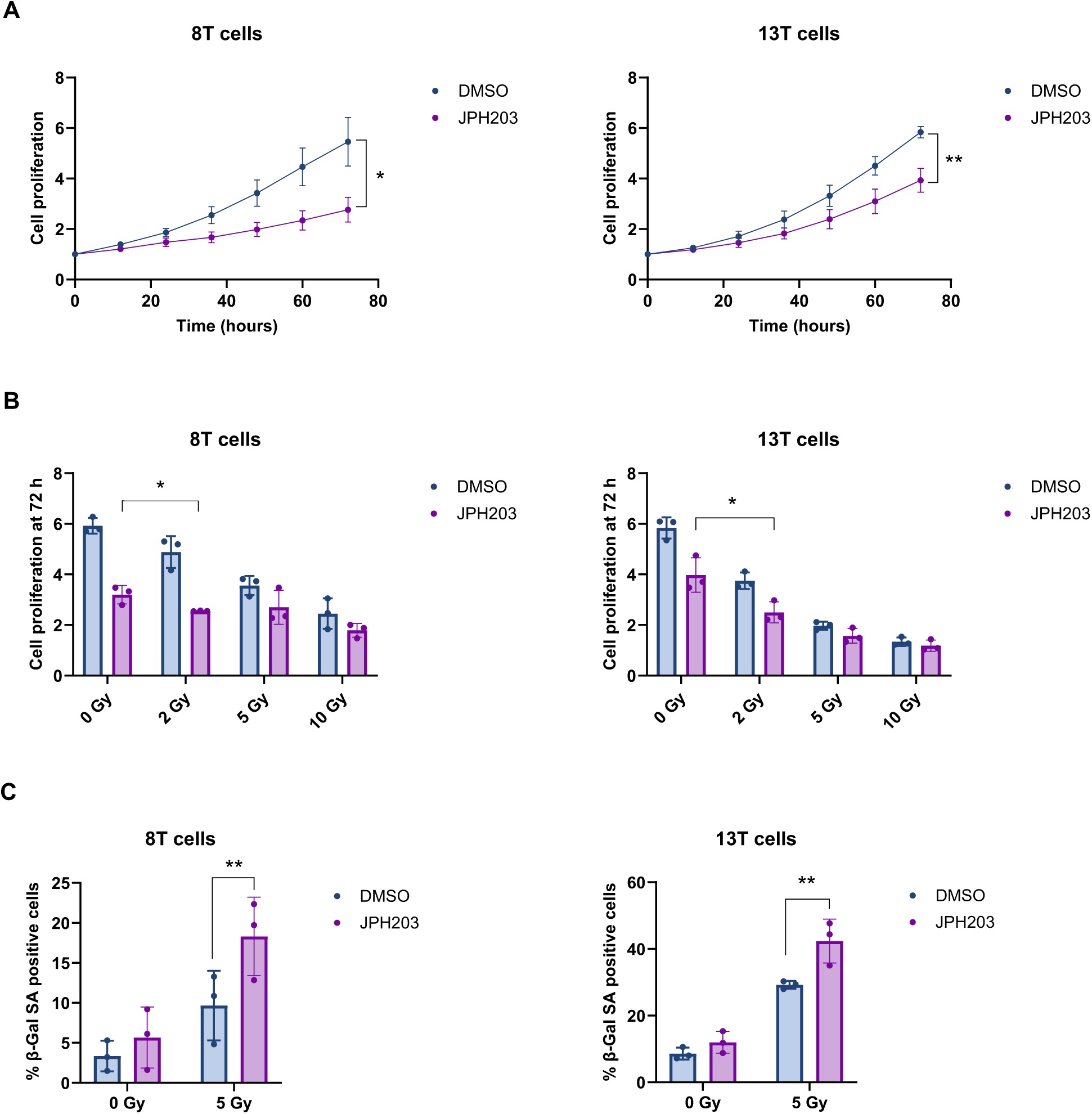
JPH203 increases senescence after radiation in MpM cells. (**A**) Cell proliferation profiles from the IncuCyte Live-Cell instrument. MpM 8T and 13T cells were treated with JPH203 (10 µM) or DMSO as a control before monitoring growth for 72 hours. Error bars represent means ± SD (n = 3 independent experiments) and significance assed by unpaired Student’s t test (* = p<0.05 and ** = p<0.01) **(B)** Cell proliferation values at 72 hours from the IncuCyte Live-Cell instrument are shown as a fold change from time 0. MpM cells were treated with JPH203 (10 µM) or DMSO one hour before X-irradiation at the indicated dose and growth was monitored for 72 hours. Error bars represent means ± SD (n = 3 independent experiments) and significance assed by unpaired Student’s t test (* = p<0.05). (**C**) Senescence associated β-gal staining assay of 8T and 13T MpM cell lines treatment with JPH203 (10 µM) or DMSO and X-irradiation with 5 Gy for 72 hours. Error bars represent means ± SD (n = 3 independent experiments) and significance assed by unpaired Student’s t test (** = p<0.01).

One drawback of chemotherapy is that administration results in systemic exposure which can result in severe side effects. Conversely, radiotherapy is more targeted and side effects are often confined to the area treated (*48*). We therefore wanted to determine if JPH203 could be used in combination with radiotherapy. To explore this possibility, we pre-treated our MpM cell lines with JPH203 and monitored their proliferation after exposure to radiation. Although JPH203 treatment or irradiation alone reduced cell proliferation, we observed a significant reduction of proliferation when combining JPH203 treatment with 2 Gy in both 8T and 13T cell lines (Fig. 6B). Interestingly, this combination treatment also triggered cellular senescence in a significant proportion of MpM cells (20% in 8T and 40% in 13T), as assessed by the number of senescence-associated β-galactosidase (SA-β-Gal) positive cells at 72 hours post-treatment (Fig. 6C, S7A), with the levels of apoptosis remaining unchanged (Fig. S7B). Although inducing senescence in tumour cells has been linked to metastasis and therapy resistance (*49*), it limits tumour cell proliferation, preventing further genomic instability and promotes the arrest of adjacent cells together with the recruitment of immune cells in vivo, potentially contributing to further tumour suppression (*50*). Moreover, several strategies aimed at eliminating senescent cells have been proposed (*49*), indicating that a combination of these treatments could increase efficacy. Therefore, these data suggest that the combination of JPH203 and radiation could potentially be beneficial as an initial anti-tumour therapy.

## DISCUSSION

RBPs play a central role in regulating gene expression and their dysregulated expression is associated with disease, in particular cancers (*8*, *9*). Numerous studies have focussed on the impact of individual RBPs in cancer, such as UNR (*51*) and PDIA6 (*52*) in Melanoma, and TRAP1 in ovarian cancer (*53*). However, few studies to date have examined global changes in the RBPome to identify new cancer drivers. The development of new technologies (*10*, *11*), including OOPS which is used in this current study, means that it is now possible to analyse global RBPome changes from relatively small amounts of material, allowing for the examination of RBPs in tumour-derived cells and tissues. For example, a recent study used OOPS to identify the RBPome of pancreatic cancer cells (*54*). As MpM has no known oncogenic drivers and is instead associated with the loss of tumour suppressor genes (*3*, *4*), we used OOPS to understand the contribution of post-transcriptional dysregulation, via changes in the RNA-bound proteomic landscape, to disease progression. Over 350 putative RBPs showed significant changes in their RNA binding capacity and the roles of these RBPs were consistent with metabolic reprogramming of MpM-derived cells, in conjunction with changes in expression of proteins related to DNA repair, replication and cell cycle, providing useful datasets for further analysis of RBPs for those studying this disease (Figure 1).

We have shown previously that translation is dysregulated in MpM (*7*) and we therefore directed our focus to RBPs that may contribute to this and could act as novel oncogenic drivers. In particular YBX3, which controls gene expression at multiple levels (*55*, *56*) including stabilising LAT1 mRNA (*31*) and promoting mRNA translation through the import of arginine, glutamine and leucine (*45*, *57*, *58*). Importantly, our data shows that YBX3 was upregulated in MpM patient TMAs, and there was a correlation between high expression of YBX3, LAT1 and patient survival, suggesting that this protein combination is associated with poor prognosis (Figure 4). Additionally, analysis of YBX3 in mouse tissue from a previous longitudinal study of MpM development (*32*) revealed that enhanced YBX3 expression was linked with tumour areas in this disease since YBX3 upregulation was observed not only in mice that developed pleural tumours tumours but also in those that developed diaphragmatic lesions. In addition, MpM has a long latency period and symptoms typically do not develop until the cancer has reached an advanced stage, delaying a diagnosis.

These findings are also consistent with our data which suggests depletion of YBX3 or LAT1 reduces MpM protein synthesis rates and cell proliferation (Figures 3 and 5). While it is important to recognise that LAT1 is also regulated by several transcription factors including c-Myc, ATF4 and YAP/TAZ (*59*), our data indicate that YBX3 post transcriptional control over LAT1 mRNA is sufficient to modify the levels of this protein in MpM (Figure 4). Although the mechanisms regulating YBX3 are beyond the scope of this study, we hypothesise its upregulation in MpM may be linked to increased E2F activity. YBX3 is a known target of the transcription factor E2F1 (Arakawa et al., 2004), and the expression of E2F targets is elevated in MpM (Mangiante et al., 2023; Robinson et al., 2015). Moreover, E2F activity is likely enhanced in MpM due to the common deletion of its negative regulator, p16/INK4A (*60–62*).

Due to the growth stimulatory effects of amino acids, targeting cellular amino acid transporters provides a promising avenue for cancer therapy (*9*, *57*). Moreover, LAT1 is highly expressed in a diverse range of human cancers including prostate (*63*), gastric (*64*), breast (*65*) and pancreatic cancers (*66*), and this expression is associated with adverse prognosis (*67*, *68*). LAT1 inhibition or knockdown has been shown to reduce the phosphorylation of mTORC1 substrates in cancer cell lines (*43*, *69*, *70*) and mouse xenograft models (*71*). In addition, the LAT1 inhibitor JPH203 suppresses proliferation of cancer cells *in vitro* and *in vivo* (*43*, *67*, *72*) and is currently under evaluation for a phase III clinical trial (*73*) following a successful phase II trial in patients with advanced refractory biliary tract cancer (*74*). In MpM, JPH203 also impairs LAT1 dependent amino acid transport and has a similar effect to YBX3 depletion downregulating mRNA translation and decreasing cell proliferation (Figure 5) in agreement with previous studies (*42*, *44*, *70*), and suggests that targeting LAT1 in MpM could be therapeutically beneficial.

There is a correlation between altered amino acid metabolism and tumour-derived cell survival after irradiation (*75*, *76*) and consistent with these data, direct inhibition of LAT1 by JPH203 treatment sensitises human lung carcinoma (A549) and pancreatic cancer (MIA Paca-2) cell lines to radiation (*77*). Our data show that LAT1 inhibition also sensitises MpM cells to radiation, reducing proliferation and inducing cell senescence. The induction of senescence by anti-cancer therapy is common (*78*), and while it can be beneficial, it is a double-edged sword. For instance, senescence has been linked to metastasis, therapy resistance, and the promotion of inflammation through the senescence-associated secretory phenotype (SASP) (*79*). However, senescence reduces cancer cell proliferation, minimises further genomic instability, supports the recruitment of immune cells, and can be targeted by the combinatorial treatment with senolytic compounds (*49*). The benefits of inducing senescence are therefore cancer or even cell type specific (*79*) and would need to be further explored in context of MpM. Nevertheless, as toxicity from JPH203 in clinical trials has been minimal (*74*, *80*), these data suggest that a combination of JPH203 and radiotherapy may represent a potentially less toxic therapy compared to the current chemotherapy regime used for MpM patients. Further studies in *in vivo* MpM models will clarify the effects of combinatorial vs monotherapy on MpM development.

There are some limitations to the approaches we have chosen. We utilised primary MpM cells because they offer a more representative pre-clinical model than commercially available cell lines that often lack key molecular features associated with the disease. However, the degree of donor-to-donor variability was not explored and may in part explain the subtle differences we observed between MpM cell lines. Additionally, we cannot exclude the possibility that YBX3 upregulation affects other YBX3-RNA complexes, potentially exerting additional proliferative or secondary functions

## METHODS

### Cell culture

Mesothelioma-derived primary cells Meso-7T and Meso-8T were isolated as described (*12*) in addition to Meso-13T(*7*), using the same methodology and displaying the same mesothelioma cell characteristics. Mesothelioma-derived primary cells were cultured in Roswell Park Memorial Institute Medium (RPMI)-1640 growth media with L-glutamine (2 mM) (Thermo 21875091) supplemented with hEGF (20 ng/ml), hydrocortisone (1 μg/ml) and 10% fetal bovine serum (FBS) at 37 °C and 5% CO_2_. Primary human mesothelial cells were purchased from Innoprot (P10770) and maintained in the same conditions as mesothelioma-derived cells for 2-3 passages. For all experiments, mesothelioma primary tumour cells and healthy mesothelial controls were plated in RPMI with L-glutamine (2 mM), 10% FBS, but with no additional growth factors and adapted at 37 °C and 5% CO2 for 18 h, before being treated as indicated. Cell proliferation was assessed using an IncuCyte S3 live-cell imaging (IncuCyte S3, Sartorius).

### Cell treatment

LAT1 was inhibited by treating cells with JPH203 (10 uM) (SelleckChem S8667). X-irradiation was performed using a CIX1 Xstrahl bench top irradiator at 160 kV and 8 mA.

### Cell transfection

Cells were transfected with siRNAs using Lipofectamine RNAiMAX reagent (Invitrogen) in six-well plates as per the manufacturer’s guidelines for 24 to 72 hours. ON-TARGETplus SMARTPool siRNAs were purchased from Horizon Discovery: YBX3 (L-015793-00), LAT1 (L-004953-01), and non-targeting control pool siRNA (D-001810-10).

### Western blot analysis

Whole-cell extracts were prepared in RIPA buffer (50 mM tris (pH 7.5), 150 mM sodium chloride, 1% Triton X-100, 0.1% SDS, 0.5% sodium deoxycholate, 1× Roche protease inhibitor cocktail, and 1× Roche PhosSTOP phosphatase inhibitor cocktail), and protein concentration was quantified using a Pierce BCA protein assay kit (Thermo Fisher Scientific). Extracts were diluted in SDS loading buffer (50 mM tris (pH 6.8), 2% SDS, 10% glycerol, 0.1% bromophenol blue, and 50 mM DTT), and total protein (20 to 30 µg) was separated using SDS–polyacrylamide gel electrophoresis. Proteins were transferred to PVDF membrane (1620177, Biorad) and incubated with the appropriate antibody at the manufacturers’ recommended dilution. Primary antibodies used were DBPA (YBX3) (#NBP1-71827, Novus), LAT1 (#5347, CST), eEF2 (#sc-166415, Santa Cruz), p-eEF2 (Thr56) (#2331, CST), eIF2α (#9722, CST), p-eIF2α (Ser51) (#109202, Abcam), p–4E-BP1 (Ser65) (#9451,CST), 4E-BP1 (9644, CST), p–p70 S6K (Thr389) (#9205, CST), p70 S6K (#2708, CST), β-actin (#A5441, Sigma), DDX3 (#8192, CST), LAT1 (#5347, CST) SLC3A2 (#HPA017980, Atlas antibodies) and β-tubulin (#2146, CST). For enhanced chemiluminescence (ECL) detection, membranes were incubated with HRP conjugated α-mouse (#7076, CST) or α-rabbit (#7074) secondary antibodies. Detection was performed using the enhanced chemiluminescence procedure (ECL Prime Western Blotting Detection Reagent, GE Healthcare #RPN2236.

### Sucrose density centrifugation

Sucrose gradients (10-50% w/v) were used to separate subpolysomal and polysomal ribosomes. These were prepared in a gradient buffer (100 mM NaCl_2_, 5 mM MgCl_2_, 15 mM tris-HCl (pH 7.5), 1 mM DTT, and cycloheximide (0.1 mg/ml)). Cells were seeded on a 15 cm plate, washed in phosphate-buffered saline (PBS)–cycloheximide (100 μg/ml), and scraped into lysis buffer (100 mM NaCl_2_, 5 mM MgCl_2_, 15 mM tris-HCl pH 7.5, 1 mM DTT, 0.2 M sucrose, 0.1 mg/ml cycloheximide, 0.5% IGEPAL, and 5 μl of RNasin per 1 ml). Lysates were incubated on ice for 3 min before pelleting cells at 1300 g for 5 min. The supernatant was layered onto the gradient and centrifuged at 38,000 rpm for 2 hours at 4°C using a Beckman Coulter ultracentrifuge. Polysomes were recorded by measuring the absorbance at 254 nm using a UA-6 UV-visible detector (Presearch Ltd.) in a gradient fractionation system (Presearch Ltd.).

### Radioisotope incorporation

For [35S]-methionine incorporation, cells were grown in 12 well plates, incubated with radiolabelled [35S]-methionine for 30 min at normal cell culture conditions. Cells were washed with PBS and lysed in RIPA buffer. Protein was precipitated using 25% TCA and protein was captured on glass fibre filter paper (GE Healthcare). Captured protein was washed with 70% ethanol and acetone before the addition of 2 ml of Ecoscint scintillation cocktail (National Diagnostics), and radioisotope incorporation was quantified using a Tri-Carb 4910TR Liquid Scintillation Counter (PerkinElmer). Each sample was carried out in triplicate, and spectral counts per minute were normalised to the total amount of protein for each sample.

### Flow cytometry

Flow cytometry data were acquired using BD LSR Fortessa (BD Biosciences). A total of 10,000 counts were acquired for each experimental condition, and all flow cytometry data were analysed with FlowJo data analysis software (v 10.9.0) (FlowJo LLC, Ashland, USA). Three independent experiments were performed. Cell death was quantified by measuring annexin V–FITC binding (488-nM laser) and Draq7 uptake in the cell (561-nM laser). Cells were collected and washed in PBS before resuspension in the annexin buffer (BD Biosciences) and incubation with annexin V and Draq7 for 20 minutes before analysis. For cell cycle analysis, cells were collected and fixed in ice-cold 70% ethanol and incubated with 1% Triton-X in PBS, 0.1 mg/ml RNase A and 20 ug/ml propidium iodide (PI) overnight.

### Senescence-associated β-galactosidase (SA β-gal) staining

Cells were plated on 6-well plates and incubated overnight. After 1-hour JPH203 pretreatment, subsequent irradiation at indicated doses and incubation for 72 hours, the cells were stained with a senescence-associated β-galactosidase staining kit (#9860, Cell Signaling Technology) according to the manufacturer’s instructions. Senescent cells were identified and counted in the light microscope EVOS XL Core (Invitrogen).

### Orthogonal organic phase separation (OOPS)

Cells were seeded in 10 cm plates, washed with cold PBS and cross-linked with UVC (400 mJ/cm^2^) irradiation prior to lysis in TRIzol™ Reagent (Thermo Fisher Scientific). The protocol was followed as described (Queiroz et al., 2019), but only one step of RNA digestion was performed with 4 μl RNase A, T1 mix (2 mg/mL of RNase A and 5,000 U/mL of RNase T1, Thermo Fisher Scientific) and incubation overnight at 37 °C, prior a final cycle of TRIzol and chloroform biphasic extraction was made and released proteins recovered from the organic phase by methanol precipitation. For western blot analysis, methanol-precipitated proteins were resuspended in 100 mM triethylammonium bicarbonate (TEAB) and 1% SDS. For mass spectrometry analysis, precipitated proteins were individually solubilized in 1% RapiGest (#186001861, Waters™) by incubation at 80°C for 10 minutes under agitation (700 rpm), and protein concentration was estimated using the Bradford protein assay (#22662, Pierce™). 100 µg of total protein was in-solution digested. For this, the protein samples were reduced with 4 mM DTT (final concentration; # R0861, Thermo Scientific™) at 60°C for 10 minutes, alkylated with 14 mM IAA (final concentration; # I3750, Sigma-Aldrich) for 30 minutes at room temperature in the dark, and quenched DTT. Trypsin digestion (1:50; sequencing-grade trypsin: protein, #V5280, Promega Corporation) was performed at 37°C for 16 hours with constant shaking (500 rpm). Peptides were quantified using the Colorimetric Peptide Assay (#23275, Thermo Scientific™) and labeled with TMTpro (#A44520, Thermo Scientific™) according to the manufacturer’s instructions. Peptides were fractionated using Reverse-phase Fractionation (#84868, Thermo Scientific™), followed by drying in a speed vacuum and storing at -20°C until use. For LC-MS/MS acquisition, peptides were resuspended in 20 µL of 0.1% TFA containing 3% acetonitrile. Peptides were analysed using an Ultimate 3000 RSLC™ nano system (Thermo Scientific™) coupled to an Orbitrap Eclipse™ mass spectrometer (Thermo Scientific™). The peptide samples were loaded onto a trapping column (Thermo Scientific™, PepMap100, C18, 300 µm x 5 mm) using partial loop injection for 3 minutes at a flow rate of 15 µL/min with 0.1% (v/v) FA in 3% acetonitrile. The sample was resolved on an analytical column (Thermo Scientific™, Easy Spray C18, 75 µm x 500 mm, 2 µm) at a flow rate of 300 nL/min using a gradient from 97% A (0.1% formic acid) to 3% B (80% acetonitrile, 0.1% formic acid) over 105 minutes, followed by an increase to 40% B over an additional 15 minutes 90% B.

LC-MS/MS data was acquired using data-dependent acquisition (DDA) program consisting of a full scan MS at a resolution of 120,000 (AGC set to 100%; 4e6 ions) with a maximum fill time of 50 ms). For TMT-MS2, fragmentation was performed using a CID activation energy of 30%. The AGC target in MS2 was set to 1E5, the maximum injection time to 50 ms, and the analyzer to ion trap with rapid detection. For TMT-MS3, fragmentation in MS2 was performed with an HCD activation energy of 55%, an AGC of 1E5, a maximum injection time of 120 ms, and the analyzer set to Orbitrap.

Raw data was imported and processed in Proteome Discoverer v2.5 (Thermo Scientific™). The raw files underwent iterative database searches using Proteome Discoverer with SequestHF and Inferis rescoring algorithms against the Homo sapiens database (updated on 24th May 2022), which contains human protein sequences from UniProt/Swiss-Prot. Common contaminant proteins (various types of human keratins, BSA, and porcine trypsin) were included in the database. Spectra identification was performed using the following parameters: MS accuracy set to 10 ppm (orbitap), MS2 accuracy of 0.5 Da (ion trap) for MS3 spectra acquired in the Orbitrap with 10 ppm, allowance for up to two missed cleavage sites, carbamidomethylation of cysteine as a fixed modification, and oxidation of methionine as a variable modification. The Percolator node was utilized for false discovery rate estimation and only rank 1 peptide identifications with high confidence (FDR < 1%) were accepted.

### RNA interactome capture (RIC)

For RIC combined with western analysis, cells were plated in 15 cm plates, washed with cold PBS and cross-linked with UVC (150 mJ/cm^2^) irradiation. Cells were harvested by scraping into lysis buffer (20 mM Tris pH 7.4, 500 mM lithium chloride (LiCl), 0.5% lithium dodecyl sulphate (LiDS), 1 mM EDTA) supplemented with 1X Complete EDTA-free protease inhibitors. Samples were passed repeatedly through a 23G needle to achieve lysis and lysates were clarified by centrifugation, DTT was added at a 5 mM final concentration. Lysates were incubated with oligo(dT) beads for one hour at room temperature with mixing. The beads were washed for 5 minutes with lysis buffer, and then washed twice for 5 minutes with 20 mM Tris pH 7.4, 1mM EDTA, 0.5 M LiCl, 0.1% LiDS, 5 mM DTT. Subsequently, the beads were washed twice for 5 minutes with 20 mM Tris pH 7.4, 0.5 M LiCl, 1 mM EDTA and twice without incubation in 20 mM Tris pH 7.4, 0.2 M LiCl and 1 mM EDTA. RNA was eluted after resuspending the beads in 20 mM Tris pH 7.4 and 1 mM EDTA and incubating the beads at 75 C for 3 minutes. RNA was digested with 2 µL of RNase A, T1 mix (2 mg/mL of RNase A and 5,000 U/mL of RNase T1, Thermo Fisher Scientific) and 125U benzonase for 4 hours. RIC samples were subjected to western analysis to investigate specific RNA-protein interactions and samples were normalised according to RNA content prior to RNA digestion.

### RIP-qPCR

Cells were seeded in a 15 cm plate, washed with cold PBS and cross-linked with UVC (150 mJ/cm^2^) irradiation. Cells were harvested by scraping into lysis buffer (50 mM Tris-HCL pH 7.4, 100 mM NaCl, 1% Igepal (NP40), 0.1% SDS, 0.5% sodium deoxycholate) supplemented with 1X Complete EDTA-free protease inhibitors and 200 U of RNaseIN (40 U/µl) per sample. After samples were passed repeatedly through a 23G needle to achieve lysis, they were incubated with 4 U of TurboDNase for 15 minutes on ice and clarified by centrifugation. Protein concentration was quantified using a Pierce BCA protein assay kit (Thermo Fisher Scientific) and samples normalised. Between 800 µg and 1 mg of protein was diluted in 1.1 ml of lysis buffer and then these lysates were incubated for 2 hours at 4°C with mixing Dynabeads Protein A (#1002D, Invitrogen) previously coupled with YBX3 antibody (#NBP1-71827, Novus) for 1 hour at room temperature with mixing. The supernatant was removed and an unbound aliquot was retained for western blot analysis. Beads were washed three times with high salt buffer (50 mM Tris-HCL pH 7.4, 1M NaCl, 1 mM EDTA, 1% Igepal (NP40), 0.1% SDS, 0.5% sodium deoxycholate) and then transferred to proteinase K buffer (10 mM Tris-HCL pH 7.4, 150 mM NaCl, 10 mM EDTA, 5 mM CaCl_2_) supplemented with SDS at a final concentration of 0.25% (w/v), DTT at 0.6 mM and 1 µl of Proteinase K (Thermofisher). Proteins were digested for 1 hour at 37 °C with mixing. For RNA extraction, samples were treated with Phenol/chloroform/isoamyl alcohol and RNA was precipitated with EtOH and NaCl. Reverse transcription was performed using SuperScript III (Thermofisher) following manufacturer’s instructions. RT-qPCR was completed using SensiFAST SYBR® Lo-ROX Kit, two-step FAST reaction in the Applied Biosystems QuantStudio 6 Flex Real-Time PCR System. The primers used were ActinB_F (5’ CTTAGTTGCGTTACACC 3’), ActinB_R (5’ ATTGTGAACTTTGGGGG 3’), SLC7A5_F (5’ ACCCTGCAGCGGAACATC 3’), SLC7A5_R (5’ CCTCCAGCATGTAGGCGTAG 3’), SLC25A6_F (5’ GCATCGTGGACTGCATTGTC 3’) and SLC25A6_R (5’ AGTAGCGAATGACGTTGGCA 3’). Fold change was calculated using the 2−ΔΔCt method.

### Proteomics data analysis

All statistical analyses and visualizations were performed using R. Peptide-spectrum matches (PSMs) from Proteome Discoverer software were used for quantification. PSMs matching contaminant proteins, multiple master proteins, PSMs with missing values in some samples, low signal:noise threshold, low intensity or high interference were excluded. PSMs intensities were used to calculate peptide intensities and then aggregated to protein level by taking the median peptide abundance. This was performed using the MSnbase R package (*81*). Log2-transformed protein abundance was centre-median normalised within each sample. Differential abundance was assessed with the Limma package from Bioconductor (*82*). The p-value was adjusted for multiple tests with the Benjamini and Hochberg method (FDR, false discovery rate). Unless otherwise indicated, a threshold of 5*10^−2^ was chosen for the adjusted P value, and a cut-off of 1.5-fold minimal change in protein abundance. Functional enrichment analysis including Gene Ontology (GO) terms and pathways from KEGG Reactome were performed using gprofiler2 R package and selecting driver terms (highlight = TRUE).

### Human TMA construction

TMA blocks containing samples from n=512 patients diagnosed with Malignant pleural mesothelioma were prepared as previously described (*7*). Each TMA block (n=12) consisted of a recipient formalin-fixed paraffin-embedded block containing 3 × 1 mm donor core donor samples per case, with 40 cases sampled per TMA block. In all, 4.5 µm sections were prepared from the finished TMA for immunostaining.

### Human TMA Immunostaining

Immunohistochemical staining was performed using multiplex immunofluorescence in a single batch using a Ventana Discovery Ultra staining platform (Roche, UK). PPolyclonal anti-YBX3 (NBP1-88049, 1:500, Novus Biologicals, UK), polyclonal anti-LAT1 (HPA052673, 1:50, Sigma Aldrich, UK) and monoclonal anti-MNF116 (M08211:350, Agilent, UK) were stained sequentially and labelled with either Omni-map anti-rabbit or anti-mouse secondary anti-sera (Roche, UK) and visualised using Opal570, Opal690 and Opal520 respectively (Akoya Bioscience, USA) and counterstained with DAPI (all Roche, UK). All immunostained slides were scanned using a Vectra3 multispectral quantitative imaging system (Akoya, USA). Whole slide images (.qptiff) were opened in Phenochart and each TMA core defined as an individual ROI. Multispectral images (.im3) were collected from each individual ROI/tissue microarray core and spectrally unmixed using InForm and tissue autofluorescence subtracted. An InForm project was established to 1) segment tissue into either ‘background glass’ (and excluded from future analysis), ‘non-tumour’ and ‘tumour’ with tumour being defined by MNF-116 expression as well as morphological clues. Individual cells were then segmented primarily using the DAPI signal and membranes / cytoplasm approximated by nominal distance from the nuclei. Using the pathologist view option in InForm, positive and negative cells were manually defined for each phenotype using multiple training images from multiple patients and including patients with sarcomatoid, epithelioid and biphasic subtypes. Following training, the InForm project was applied to all multispectral images and all output files (.txt summaries and .jpg segmented images) collected. Using R, the “tissue_seg_data_summary.txt” and “cell_seg_data.txt” output files for each TMA core were summarised to measure the areas and to quantify immunostaining intensities of each phenotype. Multiple slides from the same patient sample were consolidated by summing cell counts and averaging cell ratios (ratios for positivity / negativity for each marker) and mean marker intensities. During the data consolidation we filtered out slides with low cell count (less than 300 cells) usually representing damaged or poorly stained sections. Mean intensity and marker positivity ratio values were investigated further and were assigned to “high” and “low” categories based on the overall distribution of the values. We aimed to find a threshold visually that represents the boundary between two groups in a bimodal distribution and / or halves the samples to lower and higher halves. These high/low ratio of positive cells and mean marker intensities were later correlated with survival times. Marker intensity values (representing expression levels) were extracted from the inForm (Version 2.6) pipeline output. For both the LAT1 and YBX3 markers we used the “Entire Cell Mean (Normalized Counts, Total Weighting)” intensity values for each segmented cells and compared the expression changes of these markers in each section. The software also assigns these cells into marker positive and negative categories, we calculated the correlations on the following cell categories: “LAT1- / YBX3-“, “LAT1+ / YBX3-“, “LAT1- / YBX3+“, “LAT1+ / YBX3+“.

### Mouse tissue immunostaining

We utilised samples from our previously published MpM in vivo study (*32*). In brief, mice were injected intra-pleurally with low, occupationally-relevant doses of long fibre amosite asbestos or vehicle control for up to 20 months of age. No significant pathology was identified in the diaphragms of vehicle control-treated animals although a subset of fibre-exposed animals did develop pleural lesions or established tumours by 20 months.

A broadly similar approach to examine LAT1 and YBX3 expression was utilised in the mouse model and performed using multiplex immunofluorescence in a single batch using a Ventana Discovery Ultra staining platform (Roche, UK). Polyclonal rabbit anti-Mesothelin (PA5-79698, 1:250, Invitrogen, UK), Polyclonal rabbit anti-LAT1 (PA5-33053, 1:50, Invitrogen, polyclonal anti-YBX3 (NBP1-88049, 1:500, Novus Biologicals, UK) were sequentially labelled using Omni-map anti-rabbit secondary antibody and visualised with Opal570, Opal520 or Opal690 respectively (Akoya, USA) and DAPI (Roche, UK). Slides were scanned using a Vectra3 multispectral quantitative imaging system (Akoya Bioscience, USA).

### Amino acid analysis

Intracellular samples were prepared and extracted using a modified Bligh & Dyer (methanol, water, chloroform) method (*83*). Cells were seeded in 6 well plates per condition and washed with 0.9% NaCl solution. Intracellular metabolites were extracted by scraping after adding 200 µl of cold methanol followed by 200 µl water on ice for each well. The cell extract was transferred to an Eppendorf containing 200 µl chloroform on ice. This was incubated 20 minutes with agitation at 4 °C followed by centrifugation and 100 µl of the aqueous phase was transferred to a frozen Eppendorf tube. The aqueous phase was used for liquid chromatography - tandem mass spectrometry measurements without further conditioning. Samples were analysed as previously described (*84*, *85*) using an Agilent 1290 Infinity II chromatography system and Agilent 6470 triple quadrupole mass spectrometer. 1-2 µl of sample were injected in randomised order and analytes were separated by hydrophilic interaction liquid chromatography (HILIC) using a Waters Atlantis Premier BEH Z-HILIC Column (1.7 µm particle size, 2.1 mm X 100 mm) maintained at 40°C. A constant flow of 0.6 ml/min was used, starting at 15% buffer A (50% acetonitrile in water, 10 mM ammonium formate, 0.176% formic acid) and 85% Buffer B (95:5:5 acetonitrile:methanol:water, 10 mM ammonium formate, 0.176% formic acid). Initial conditions were kept constant for 3 min, followed by ramping to 5% B over 7 min, which was kept constant for 1 min before returning to initial conditions with a total analysis time of 12 min. The mass spectrometer was operated in dynamic multiple reaction monitoring (dMRM) mode. MS parameters were as follows: Gas temp:325°C, Gas flow:10 l/min, Nebulizer:40psi, Sheath gas temp:350°C, Sheath gas flow:11 l/min, Capillary (positive):3500V, Capillary (negative):3500V, Nozzle voltage (positive):1000V, Nozzle voltage (negative):1000V, Cycle time: 500ms. Data were analysed using MassHunter Workstation Quantitative Analysis for QQQ v10.1. Chromatograms were integrated using the ‘Agile2’ or ‘Spectrum Summation’ algorithms and peak areas were converted to concentrations using standard curves obtained from serially diluted analytical standards. The data were then processed further using custom R scripts, including probabilistic quotient normalisation (*86*). A two-way ANOVA with Tukey’s multiple comparison test was performed to compare the three time points of JPH203 treatment on each amino acid.

### Statistical analysis

All statistical analyses were performed using R or GraphPad Prism 10. Statistical tests were chosen based on normality, variance, variables and study design. Details are specified in the figure legends.

## Acknowledgments

We thank the Histopathology, Mass Spectrometry and Bioinformatics facility of MRC Toxicology Unit and the CRUK Scotland Institute Histology Facility for their support throughout this study. We would like to thank M.Stoneley for critical reading of the manuscript.

## Author contributions

Conceptualization: A.E.W., M.M., and A.R. Methodology: A.R., R.F.H., M.S., C.FI., I.P., S.K., and D.F.A. Bioinformatics and data analysis: A.R., M.S., L.K., I.P., and R.G. Investigation: A.R., R.F.H., A.C., M.S., C.FI., C.FR., G.M., and S.K. Visualization: A.R., M.S., and R.F.H. Supervision: A.E.W., M.M., A.R., J.P.C.L.Q., K.P., and D.F.A. Writing—original draft: A.R., A.E.W., and R.F.H. Writing—review & editing: A.E.W., A.R., R.F.H., M.M., K.P., D.F.A., J.P.C.L.Q., and M.M.

## Data availability

The mass spectrometry OOPS proteomic raw data generated in this study have been deposited in the ProteomeXchange Consortium via the PRIDE (*87*) partner repository and are accessible with the dataset identifier PXD054230. Processed data is available in Tables S1-4.

---

**Fig. S1.**
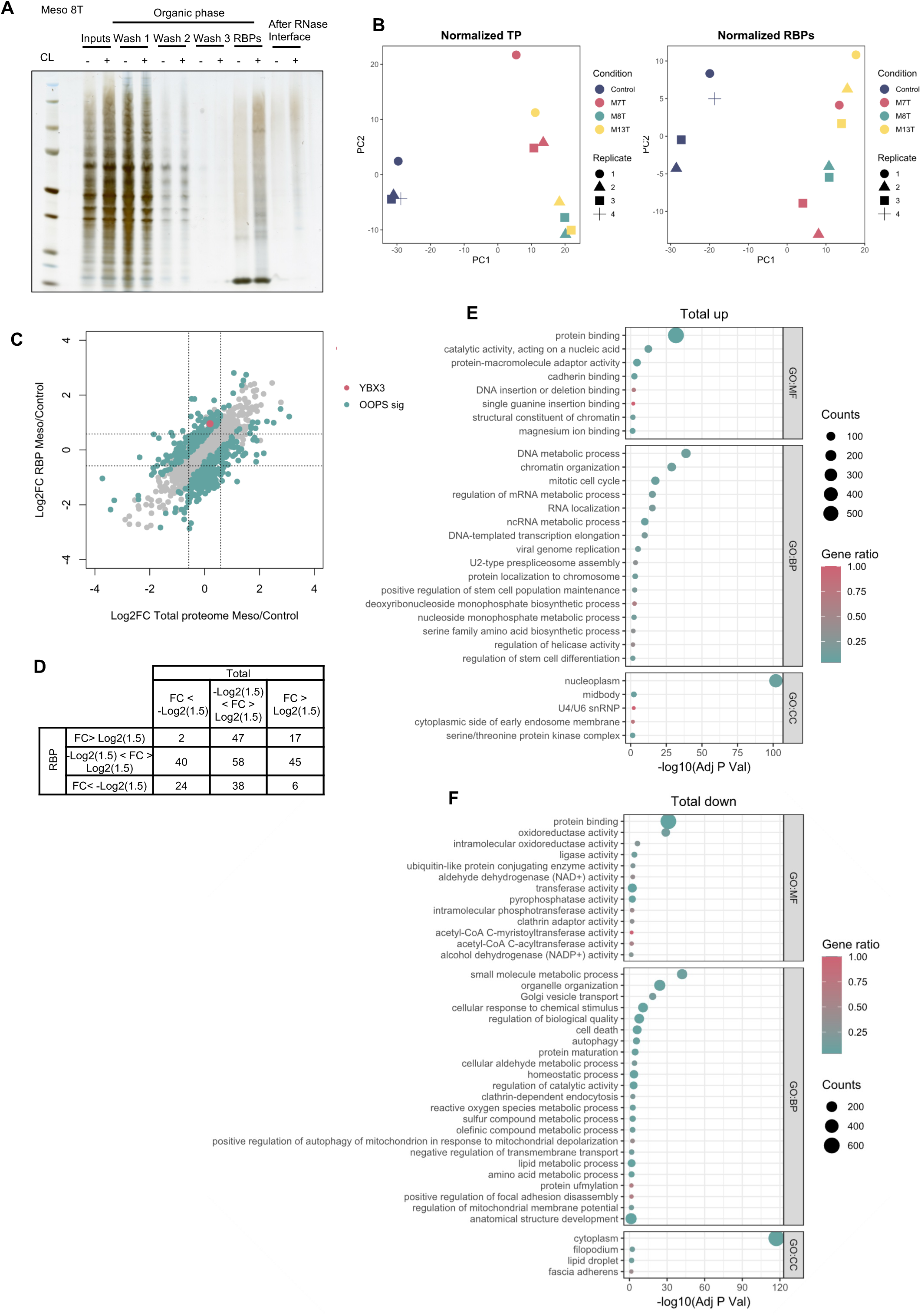
OOPS quality controls and total proteome results are consistent with a tumour proliferation phenotype. (**A**) Image of a silver-stained polyacrylamide gel showing organic phase samples from each step of the OOPS protocol with or without crosslinking (+/- CL). (**B**) PCA plots showing control and MpM samples grouping along its first component in total proteome samples (TP) and interface samples (RBPs). (**C**) Scatter plot comparing fold change from total and RBP samples in MpM cell lines against control cells. The dotted lines correspond to 1.5-fold differences. (**D**) Contingency table from OOPS significant changes represented in (C). (**E** and **F**) Dot plots depicting GO functional enrichment analysis of total proteome showing an increase (E) or decrease (F) in expression.

**Fig. S2.**
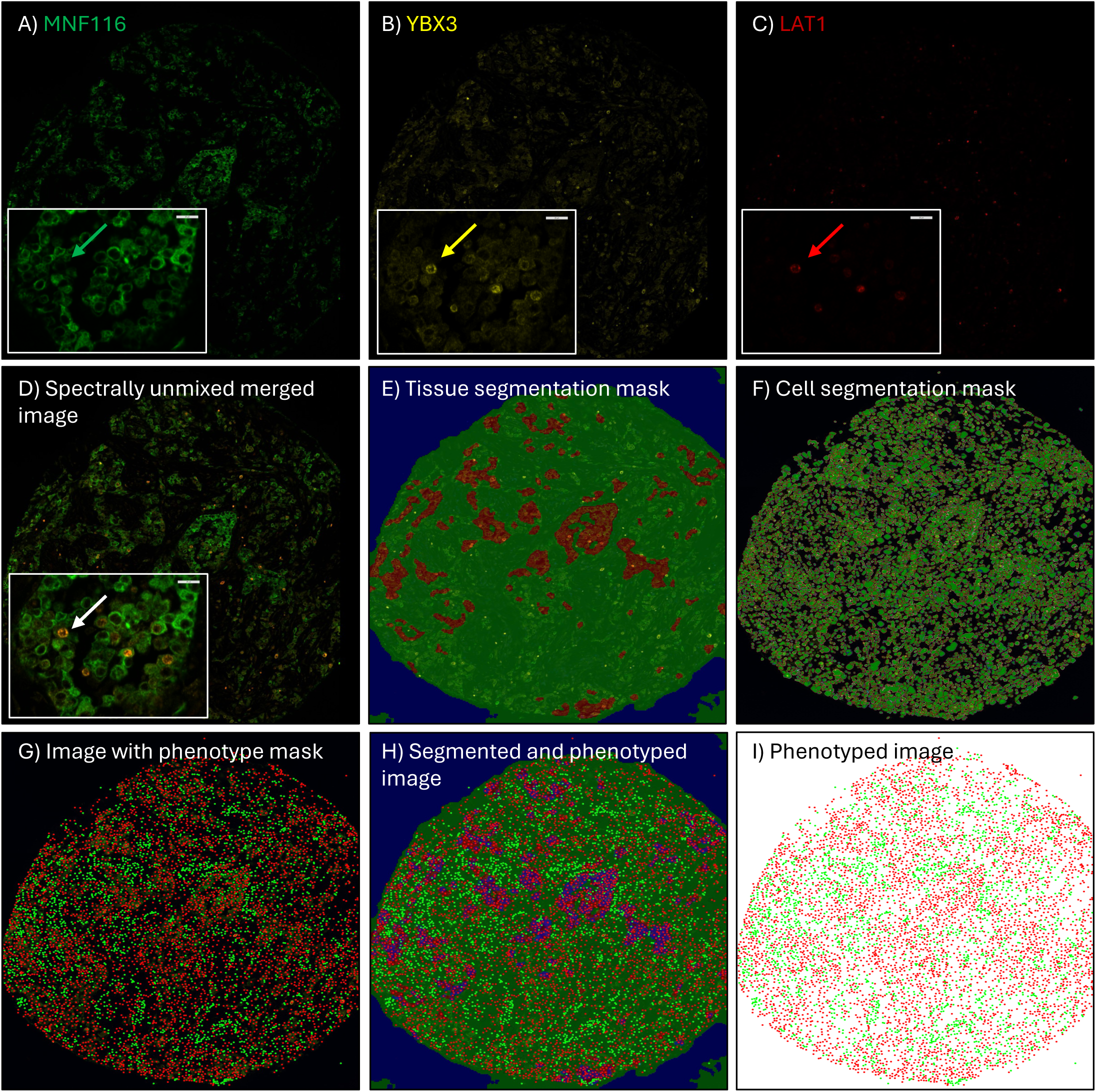
Staining of human MpM TMAs. Human TMA sections were immunostained with **(A)** anti-Human MNF116 (to aid identification of cytokeratin-rich tumour), **(B)** anti-human YBX3, **(C)** LAT1 and counterstained with DAPI. InForm software package was used to spectrally unmix individual images **(D)** to correct for spectral overlap of neighbouring fluorophores. MNF116 and morphological clues were used to train an algorithm **(E)** to segment tumour (red), stroma (green) from background glass (blue). Individual cell nuclei were identified with DAPI **(F)** and pixel intensities for each fluorophore were used to assess and quantify each cell phenotype within the tumour region **(G-I)**. This process was repeated to measure LAT1 and YBX3 positive cells.

**Fig. S3.**
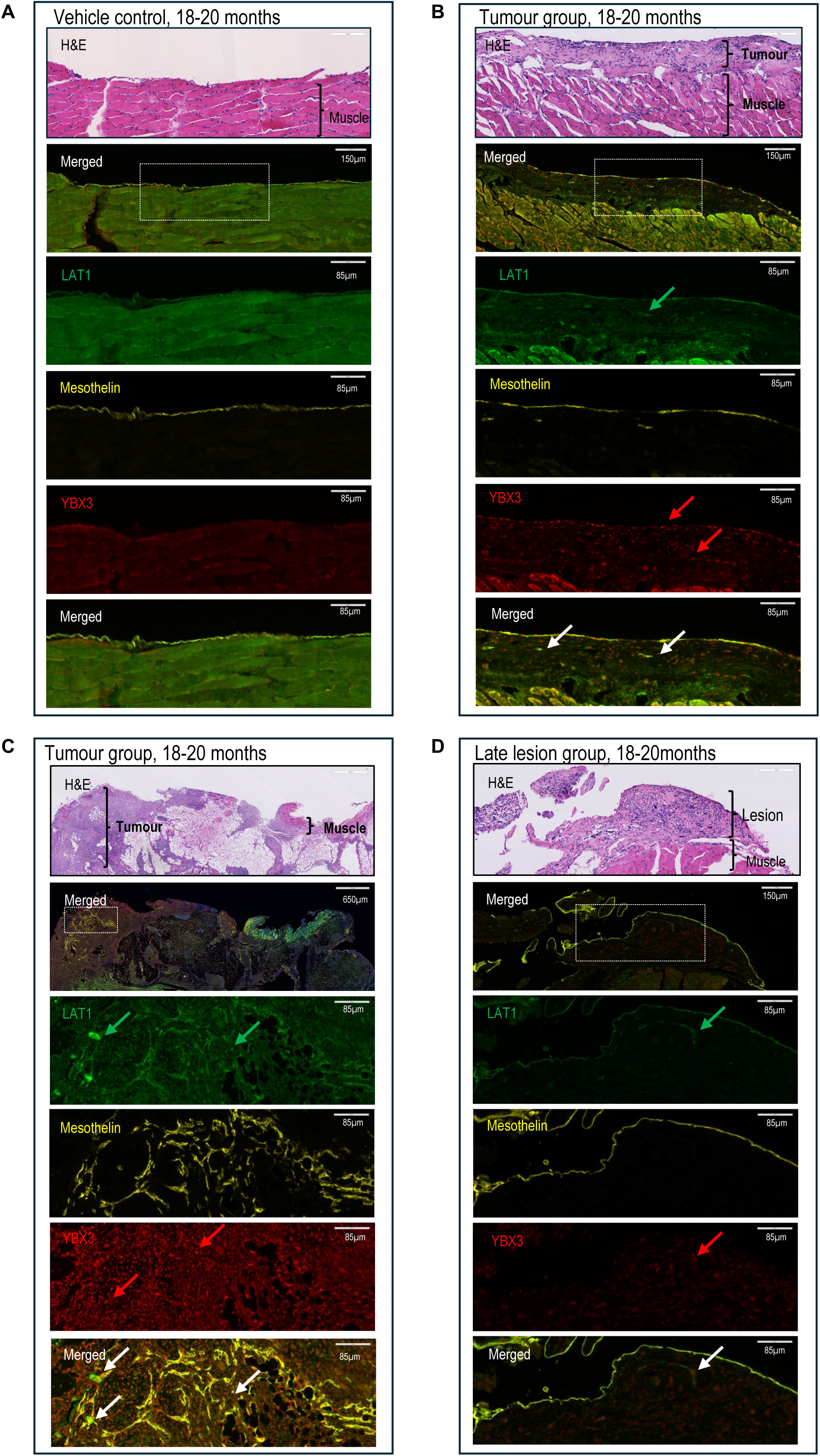
YBX3 expression is enhanced in MpM tumours. Diaphragm sections from vehicle control mice **(A)**, and from mice exposed to asbestos that developed either established tumours **(B and C)** or diaphragmatic lesions **(D)** were immunostained with LAT1, Mesothelin, YBX3 and DAPI.

**Fig. S4.**
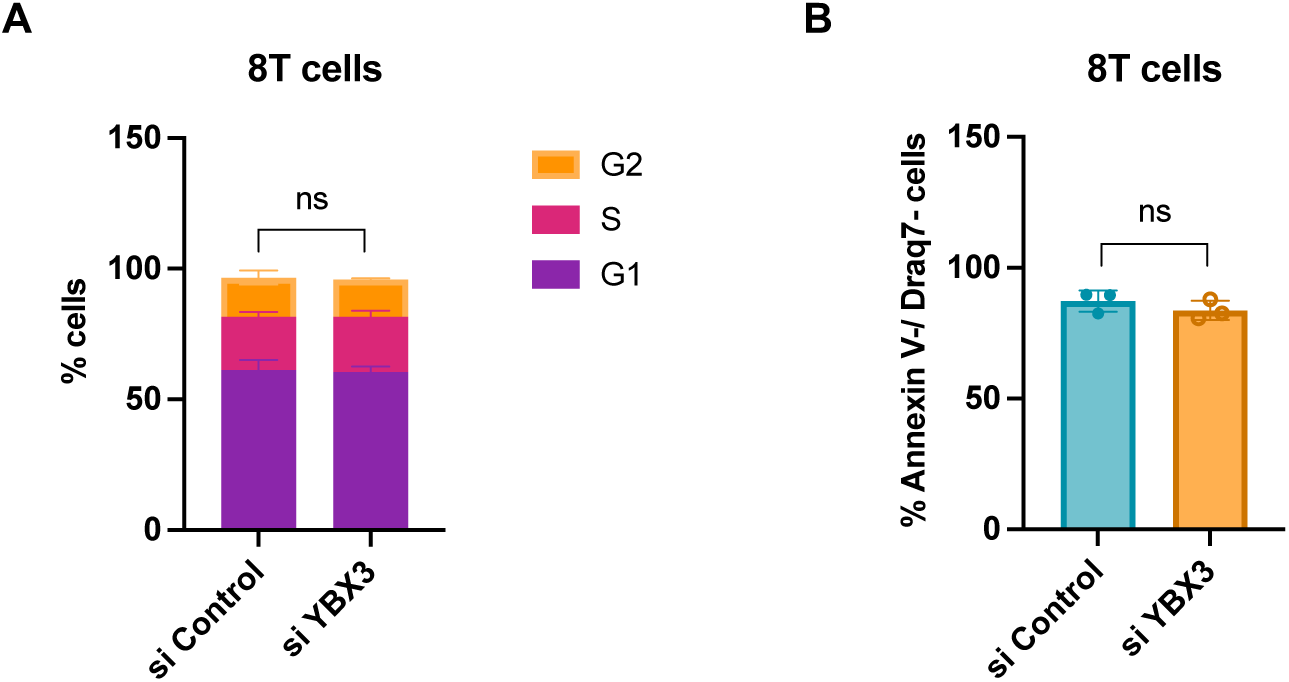
YBX3 depletion does not affect cell cycle or result in cell death. (**A**) Quantification of cell cycle state using propidium iodide (PI) staining in 8T cells treated with siRNA specific for YBX3 or non-targeting control for 72 hours. Error bars represent means ± SD (n = 3 independent experiments). (**B**) Quantification of cell death using annexin V–FITC and Draq7 staining of 8T cells treated with siRNA specific for YBX3 or non-targeting control for 72 hours. Error bars represent means ± SD (n = 3 independent experiments).

**Fig. S5.**
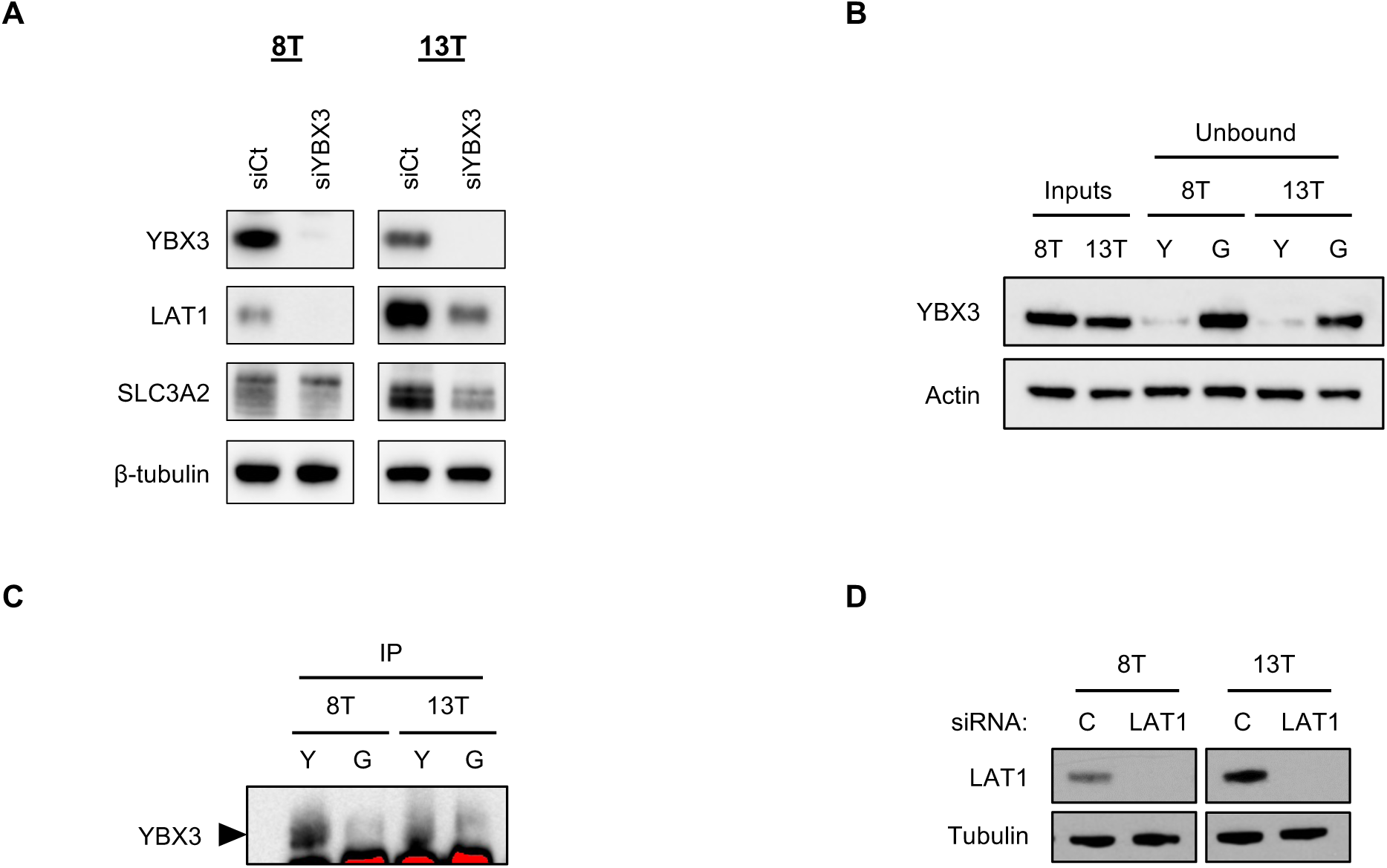
YBX3 binds LAT1 mRNA in MpM cell lines. (**A**) Representative western blots (three independent experiments) from 8T and 13T MpM cell lines comparing LAT1 and SLC3A2 expression after treatment with siRNA specific for YBX3 or non-targeting control for 72 hours. (**B**) Representative western blots (two independent experiments) of cell lysates from 8T and 13T MpM cell lines before (Inputs) and after immunoprecipitation (unbound) with beads coupled to YBX3 antibody (Y) or IgG control (G). (**C**) Representative western blots (two independent experiments) showing immunoprecipitation of YBX3 in MpM cell lines 8T and 13T, in parallel to Figure 4B. **(D)** Representative western blot of LAT1 protein expression levels confirming the knockdown efficiency at 72 hours (in parallel to Figure 4C).

**Fig. S6.**
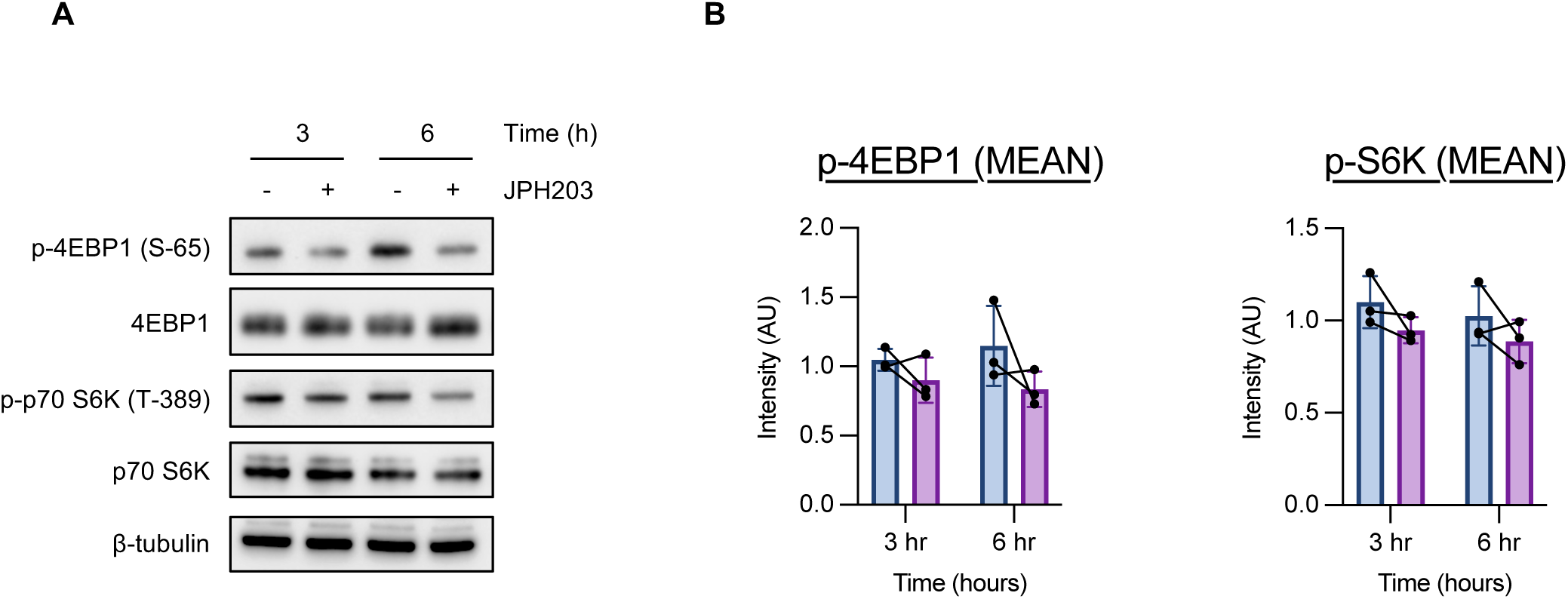
mTOR signalling in MpM 8T following JPH203 treatment. (**A**) 13T cells were treated JPH203 (10 uM) for the indicated times and analysed by immunoblotting with the indicated antibodies. **(B)** Quantification of 4EBP1 (S-65) and p70 S6K (T-389) phosphorylation from (A). Error bars represent means ± SD (n = 3 independent experiments).

**Fig. S7.**
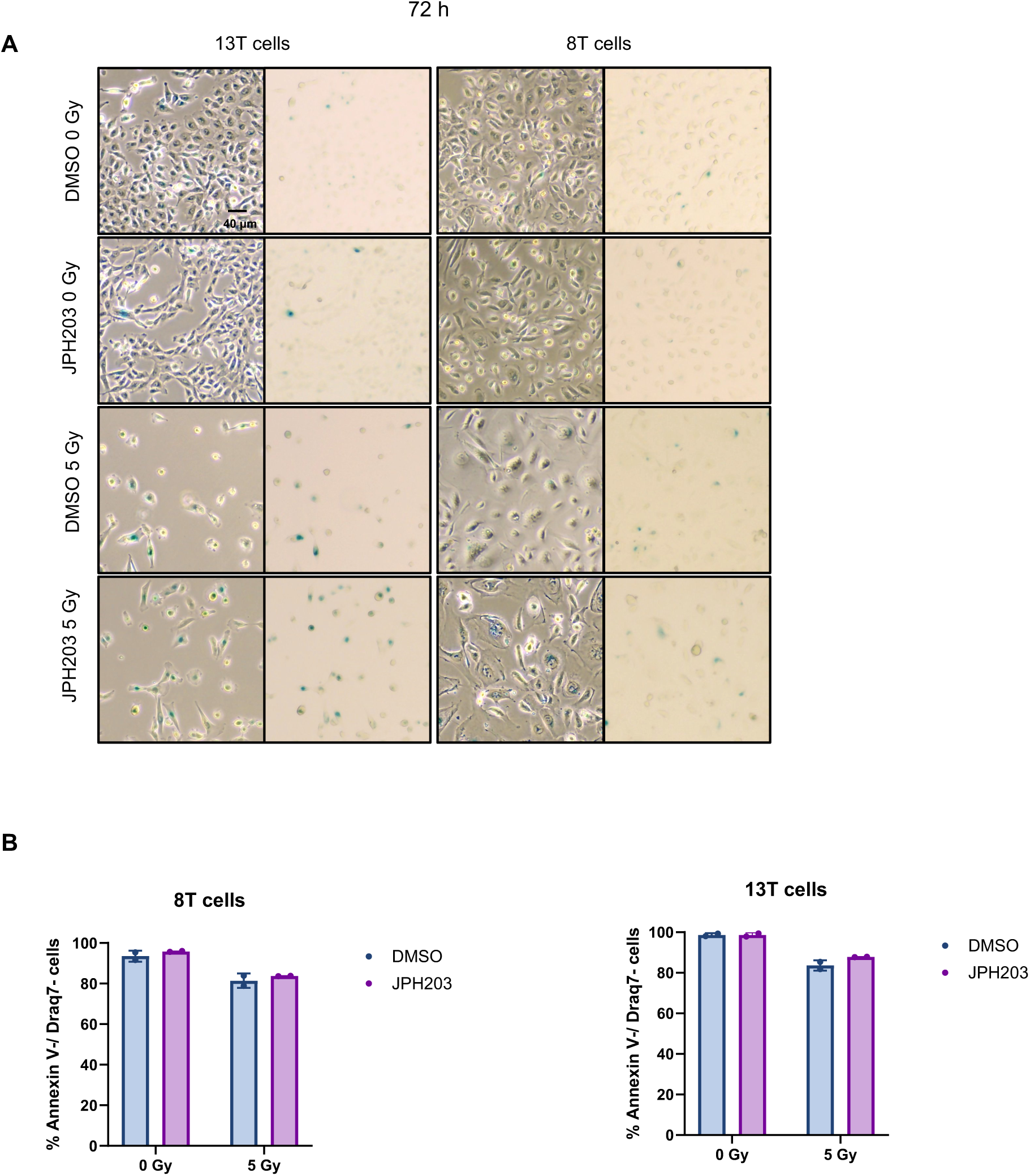
MpM senescent cells after JPH203 and X-irradiation treatment. (**A**) Representative images of SA β-gal positive cells after JPH203 or DMSO and X-irradiation (in parallel to Figure 6C). Scale bar = 40 µM. (**B**) Quantification of cell death using annexin V–FITC and Draq7 staining of 8T and 13T MpM cell lines treated with JPH203 (10 µM) or DMSO and X-irradiation with 5 Gy for 5 days. Error bars represent means ± SD (n = 2 independent experiments) with individual data points.

